# Delayed forebrain excitatory and inhibitory neurogenesis in STRADA-related megalencephaly via mTOR hyperactivity

**DOI:** 10.1101/2025.05.13.653911

**Authors:** Tong Pan, Grace Lin, Xuan Li, Debora VanHeyningen, John Clay Walker, Sahej Kohli, Aiswarya Saravanan, Amrita Kondur, Daniel C. Jaklic, Saul Pantoja-Gutierrez, Shivanshi Vaid, Julie Sturza, Ken Inoki, Tomozumi Imamichi, Weizhong Chang, Louis T Dang

## Abstract

Biallelic pathogenic variants in *STRADA*, an upstream regulator of the mechanistic target of rapamycin (mTOR) pathway, result in megalencephaly, drug-resistant epilepsy, and severe intellectual disability. This study explores how mTOR pathway hyperactivity alters cell fate specification in dorsal and ventral forebrain development using *STRADA* knock-out human stem cell derived brain organoids. In both dorsal and ventral forebrain *STRADA* knock-out organoids, neurogenesis is delayed, with a predilection for progenitor renewal and proliferation and an increase in outer radial glia. Ventrally, interneuron subtypes shift to an increase in neuropeptide-Y expressing cells. Inhibition of the mTOR pathway with rapamycin results in rescue for most phenotypes. When mTOR pathway variants are present in all cells of the developing brain, overproduction of interneurons and altered interneuron cell fate may underlie mechanisms of megalencephaly, epilepsy, and cognitive impairment. Our findings suggest mTOR inhibition during fetal brain development as a potential therapeutic strategy in *STRADA* deficiency.

## INTRODUCTION

Pathogenic variants in genes that regulate the mTOR signaling pathway, including *TSC1, TSC2, PTEN, PI3K, AKT, STRADA*, and *DEPDC5*, give rise to a spectrum of neurodevelopmental disorders collectively known as mTORopathies.^1^ These conditions share common manifestations, including brain malformations, epilepsy, and neuropsychiatric changes.^2,3^ Cortical malformations in mTORopathies can be either focal, as in focal cortical dysplasia (FCD) and tuberous sclerosis complex (TSC), or diffuse,^4^ as seen in polyhydramnios, megalencephaly, and symptomatic epilepsy (PMSE) syndrome. PMSE, a rare mTORopathy most frequently seen in the Old Order Mennonite population, is caused by inherited biallelic loss-of-function variants in the *STRADA* gene.^5^ A single post-mortem neuropathological study of a 7-month-old infant with PMSE revealed hallmark features of mTOR complex 1 (mTORC1) hyperactivation, including neuronal cytomegaly, increased phospho-S6 (pS6) expression, white matter vacuolization and astrocytosis.^6^ STRADA, a pseudokinase and upstream regulator of mTORC1 signaling,^7^ normally complexes with LKB1 and MO25, which activates LKB1 kinase activity.^8,9^ LKB1, in turn, phosphorylates and activates adenosine monophosphate-activated kinase (AMPK), which inhibits mTORC1 through the phosphorylation of Raptor, WDR24, and TSC2.^10,11^ TSC2 forms a tumor suppressor complex with TSC1 and TBC1D7,^12^ in which TSC2 acts as a GTPase activating protein toward Rheb, an essential activator of mTORC1.^13^ Loss of function variants in either TSC1 or TSC2 lead to tuberous sclerosis complex (TSC), a genetic disorder with tumor formation, intellectual disability, and epilepsy.^14^ Loss of STRADA leads to the disruption of TSC1/TSC2 repression, resulting in constitutive mTORC1 activation. Unlike autosomal dominant mTORopathies such as TSC,^15–17^ pathogenesis in PMSE does not require a second genetic hit.

Human induced pluripotent stem cell (iPSC) and mouse models have demonstrated that STRADA plays a critical role in regulating mTORC1 signaling during corticogenesis. Depletion of STRADA in mouse neural progenitors via short hairpin RNA (shRNA) knockdown led to abnormal cortical lamination and impaired neural stem cell migration, secondary to mTORC1 hyperactivation.^18,19^ Germline *Strada* knockout (KO) mice exhibited ectopic late-born Cux1-positive neurons in the intermediate zone at postnatal day (P) 0 and in the deep white matter at P9.^19^ Our prior work using human induced pluripotent stem cells (iPSCs) from PMSE patients demonstrated hallmark findings of mTORopathies: 2-D cortical-like neurons had increased pS6 and cytomegaly,^20^ and 3-D dorsally-directed human cerebral organoids (dhCOs) had increased pS6, increased phosphorylated 4EBP1, and organoid size.^21^ Additionally, dhCOs demonstrated delayed neurogenesis and an expanded outer radial glia (oRG) population, a later-born neural progenitor population that is greatly expanded in human cortical development. ORGs contribute to the increased expansion and diversity of upper layer excitatory neurons present in primates compared to other organisms such as rodents.^22^. These human-derived models offer strong construct validity by paralleling patient genetics and capturing signaling abnormalities from the earliest stages of neurodevelopment. Altogether, these results support a conserved role for STRADA in neural development, but the impact of STRADA loss on ventral forebrain development and inhibitory interneuron specification remains unexplored, despite growing evidence implicating interneuron dysfunction and excitation-inhibition imbalance in the pathophysiology of mTORopathies.^23–27^

To investigate the neurodevelopmental consequences of STRADA loss, we generated *STRADA* KO iPSC lines and differentiated them into dhCOs and ventral cortical organoids (vhCOs). Both dorsal and ventral KO organoids displayed increased size and mTORC1 hyperactivity. Delayed differentiation and increased stem cell fate was common among dorsal and ventral KO organoids. Additionally, interneuron subtype composition of ventral organoids was altered by STRADA loss. Pharmacological inhibition of mTORC1 with rapamycin rescued most of these phenotypes, supporting the role of mTORC1 hyperactivity in *STRADA*-related neurodevelopmental abnormalities. This study expands current knowledge of the role of mTOR signaling in brain development by uncovering its role in interneuron development, using patient-relevant, human-derived models.

## RESULTS

### STRADA knockout forebrain organoids display increased growth and mTOR hyperactivity

We used CRISPR/Cas9 genome editing to generate *STRADA* loss of function KO lines with compound heterozygous truncating variants and isogenic controls (Fig. S1A-1B). Cell lines were verified for pluripotency marker expression (Fig. S1C). There were no significant karyotypic abnormalities (data not shown). Two STRADA mutant lines (M1, M2) and two wildtype (WT) isogenic control lines (C1, C2) were used for all subsequent experiments. qPCR results from day 14 and 35 showed an expected depletion of *STRADA* mRNA in *STRADA* KO organoids (Fig. S1D) and significantly decreased STRADA protein in dhCOs and vhCOs for both M1 and M2 compared to C1 and C2 at day 77 (*p*<0.01, Fig. S1E-F). The expression of STRADA protein expression had an increasing trend from 0 to 77 days in dhCOs or vhCOs from C1, suggesting that STRADA is present in neural stem cells and increases with neurogenesis (Fig. S1G-H).

We adapted a directed differentiation protocol to generate dorsally and ventrally patterned cortical organoids (dhCOs and vhCOs) in parallel (Fig. S2).^28^ To assess size and growth pattern of organoids in response to STRADA loss, dhCOs and vhCOs from all four cell lines were differentiated, and organoid size was quantified from day 5 to day 35 (Fig. 1A, B). STRADA loss led to increased organoid size in both dhCOs and vhCOs when analyzed using mixed models of individual growth. The STRADA KO organoids were larger than WT with differences starting at day 13 (β=0.18, 95% CI [0.09–0.27], p <0.01 for dorsal; β=0.09, 95% CI [0.04–0.14], *p*<0.01 for ventral) and becoming more pronounced at day 35 (β=0.72, 95% CI [0.4–1.04], *p*<0.01 for dorsal; β=0.42, 95% CI [0.20–0.64], *p*<0.01 for ventral). Chronic rapamycin treatment, initiated at day 12 and maintained until organoid harvest, significantly reduced organoid size in STRADA KO (*p*<0.0001). Taken together, these results indicate that STRADA loss drives mTOR-dependent organoid overgrowth, consistent with the megalencephaly phenotype seen in PMSE patients.

**Figure 1.**
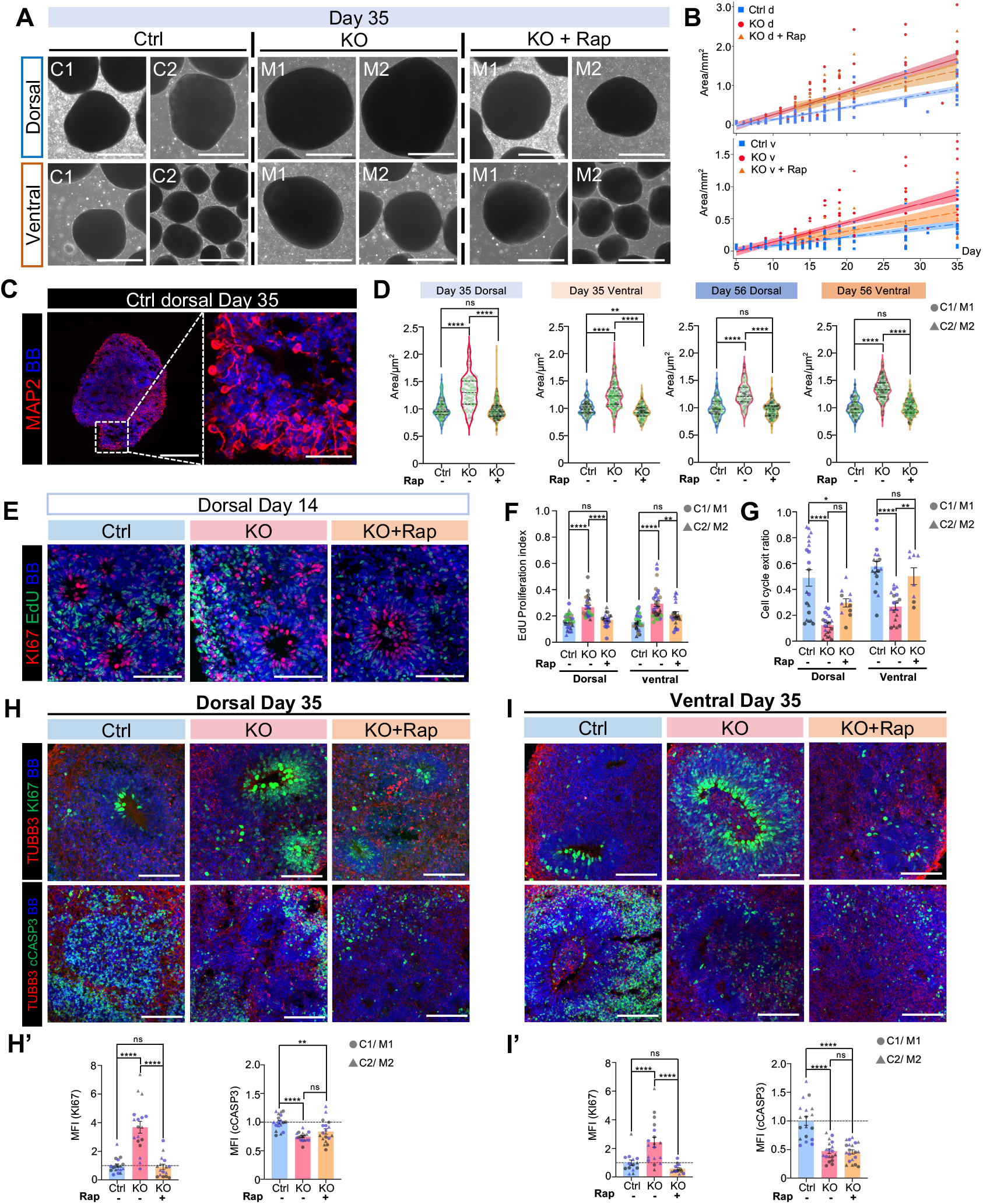
STRADA KO dhCOs and vhCOs exhibit enhanced growth and cell proliferation. **(A)** Bright field images of day 35 C1, C2, M1, and M2 dhCOs and vhCOs with vehicle or chronic rapamycin treatment. **(B)** Growth curves and statistical analysis of control and STRADA KO organoid sizes from day 5 to day 35 of differentiation with vehicle or chronic rapamycin treatment initiated at day 12. (**C**) Neuronal size in control dhCOs at day 35, measured as MAP2-outlined area. (**D)** Quantification of cell sizes for day 35 and 56 organoids. **(E)** Sample images of EdU labeling for control, KO, and KO treated with rapamycin starting day 12, harvested 2 hours after EdU pulse at day 14. **(F)** Proliferation index (PI) for panel E, calculated as EdU+ cells over all bisbenzimide (BB) cells, with organoids labeled at day 14 and harvested 2 hours post-labeling. (**G)** Cell cycle exit ratio for organoids labeled with EdU at day 14 and harvested 24 hours post-labeling; calculated as EdU+ KI67-cells over all EdU+ cells. (**H)** Immunostaining of KI67 and cleaved caspase 3 (cCASP3) in day 35 dhCOs, treated with vehicle or rapamycin. (**H’)** Quantification of results from H. (**I)** Immunostaining of KI67 and cCASP3 in day 35 vhCOs treated with vehicle or rapamycin. (**I’)** Quantification of results from I. Quantitation in H’-I’ are mean fluorescence intensity (MFI) within each organoid, with each dot representing one organoid from distinct batches of differentiations (color-coded). Data are presented as mean ± SEM, with statistical significance via one-way ANOVA. \**p*<0.05, \*\**p*<0.01, \*\*\**p*<0.001, \*\*\*\**p*<0.0001. Scale bars, A-1 mm, C-300 µm (50 µm for zoomed-in image), E/H/I-100 µm.

To investigate the mechanisms underlying increased organoid size caused by STRADA loss, we analyzed cell size, proliferation, and survival in dhCOs and vhCOs. At day 35 and 56, when neurogenesis is ongoing, STRADA KO organoids exhibited cytomegaly compared to controls, as measured by neuronal size outlined with neuronal marker MAP2 (Fig. 1C, D). Cytomegaly was rescued by chronic rapamycin treatment. To examine early neurogenesis, day 14 organoids were treated with EdU for 2 hours to label actively dividing cells and then harvested immediately thereafter or 24 hours later (Fig. S1). Both dorsal and ventral STRADA KO organoids showed a significant increase in cell proliferation index at 2 hours (Fig. 1E-F) and a decrease in cell cycle exit ratio (EdU+/Ki67-divided by all Edu+ cells) at 24 hours (Fig. 1G). Rapamycin-treated STRADA KO dhCOs and vhCOs demonstrated similar proliferation dynamics to controls (Fig. 1E-G). Additionally, day 35 organoids stained for KI67 (Fig. 1H, H’) and cleaved caspase 3 (cCASP3, Fig. 1I, I’) revealed increased cell proliferation and decreased cell death in STRADA KO organoids. Rapamycin treatment rescued the phenotype of increased proliferation but not the reduction in cell death. In all, these findings indicate that STRADA loss drives early organoid overgrowth primarily through mTORC1-dependent enhancement in cell growth and proliferation.

To visualize mTORC1 hyperactivity in STRADA KO organoids, protein levels of the downstream effectors P-S6 and P-4EBP1 were examined in day 35 organoids. Western blotting revealed a consistent increase in P-S6 and P-4EBP1 in KO dhCOs and vhCOs compared to control (Fig. 2A-D). Immunostaining showed similar results, with significantly higher P-S6 and P-4EBP1 signal in STRADA KO dhCOs and vhCOs (Fig. 2E-H). Rapamycin treatment significantly restored P-S6 and P-4EBP1 to levels comparable to control in STRADA KO dhCOs and vhCOs (Fig. 2).

**Figure 2.**
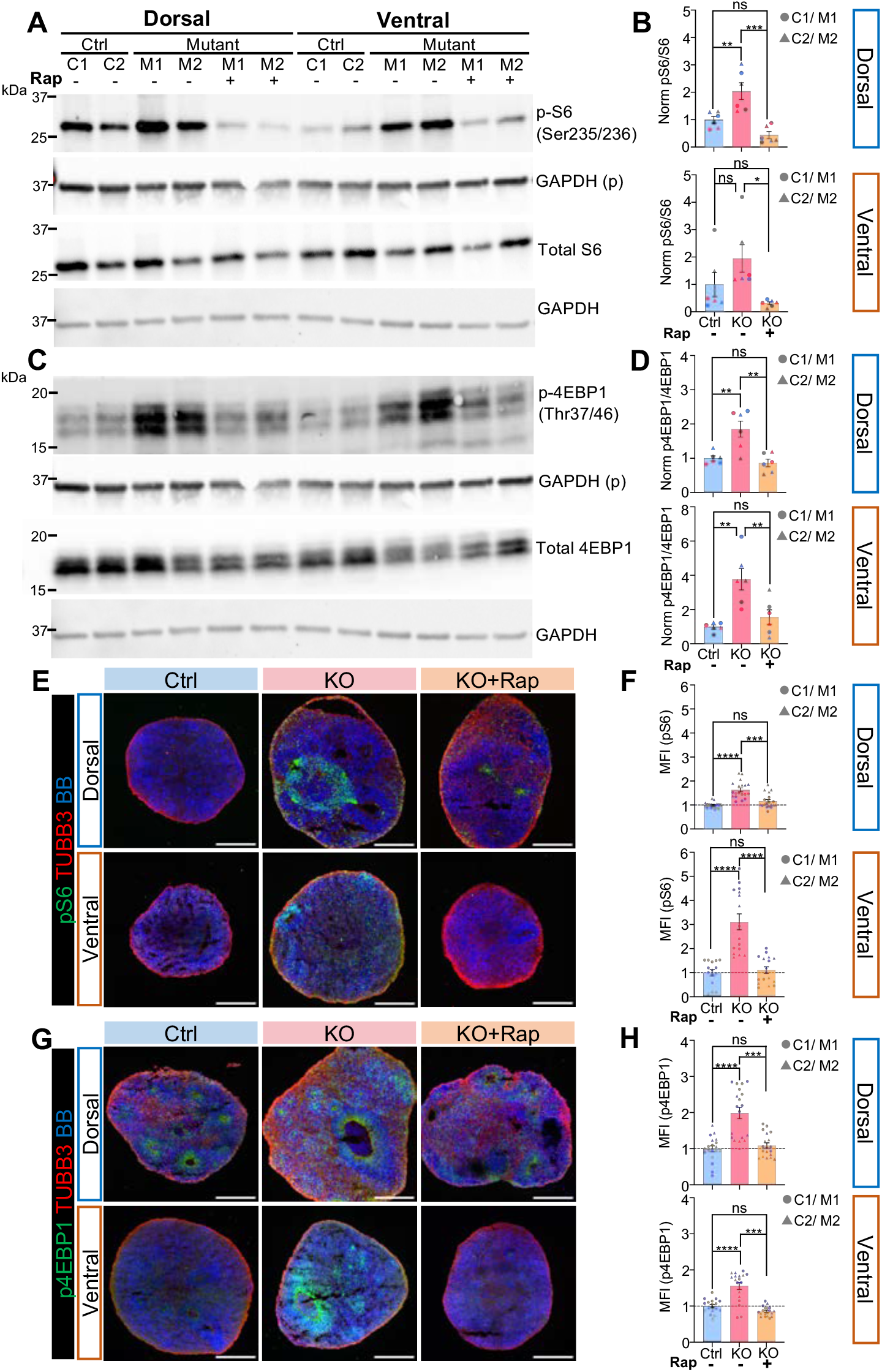
STRADA KO dhCOs and vhCOs show elevated mTORC1 activity. **(A)** Western blotting of day 35 control and STRADA KO dhCOs and vhCOs, treated with vehicle or rapamycin from day 12, for the mTOR effector P-S6 and total S6 protein, normalized to the loading control GAPDH. **(B)** Quantification of results in A, with P-S6 normalized to total S6 and each dot representing a biological replicate. **(C)** Western blotting for P-4EBP1 and total 4EBP1 proteins under the same conditions as panel A, normalized to GAPDH. **(D)** Quantification of results in C, with P-4EBP1 normalized to total 4EBP1. (**E-F)** Immunostaining images (E) and quantitation (F) for P-S6, co-stained with neuronal marker TUBB3, in control and STRADA KO dhCOs and vhCOs, treated with vehicle or rapamycin. (**G-H)** Immunostaining images (G) and quantitation (H) for P-4EBP1 under the same conditions. Each shape represents one organoid, compiled from separate batches of differentiation (color-coded). For all quantifications mean ± SEM is displayed, with Kruskal-Wallis test. \**p*<0.05, \*\**p*<0.01, \*\*\**p*<0.001, \*\*\*\**p*<0.0001. Scale bars, E/G-300 µm. Uncropped western blots are available in Fig. S7.

### Dorsal and ventral organoids undergo proper forebrain regionalization, with altered neurogenesis dynamics in response to STRADA loss

To validate forebrain regionalization, we examined dhCOs and vhCOs for region-specific markers at key developmental stages: day 35 (newborn neuron production), day 56 (ongoing neurogenesis with multiple terminally differentiated neuronal subtypes being born), and day 77 (mature neuron specification). At days 35 and 56, dhCOs expressed the dorsal progenitor marker PAX6, while vhCOs robustly expressed the ventral progenitor marker NKX2.1, confirming proper regional identity (Fig. 3A–B). STRADA KO organoids exhibited increased PAX6 and NKX2.1 expression compared to controls at both timepoints, consistent with expanded neural progenitor pools. All organoids, regardless of genotype and regionalization, expressed the forebrain marker FOXG1 at day 56 (Fig. S3), supporting forebrain cell fate specification. Rapamycin normalized PAX6 and NKX2.1 levels in STRADA KO organoids to control levels, suggesting that mTOR hyperactivity drives progenitor expansion. Further analysis at day 77 demonstrated strong GAD67 expression in vhCOs but minimal signal in dhCOs, confirming successful ventralization and interneuron production (Fig. 3C–D). Notably, STRADA KO vhCOs exhibited reduced GAD67 levels compared to controls, suggesting disrupted GABAergic interneuron maturation.

**Figure 3.**
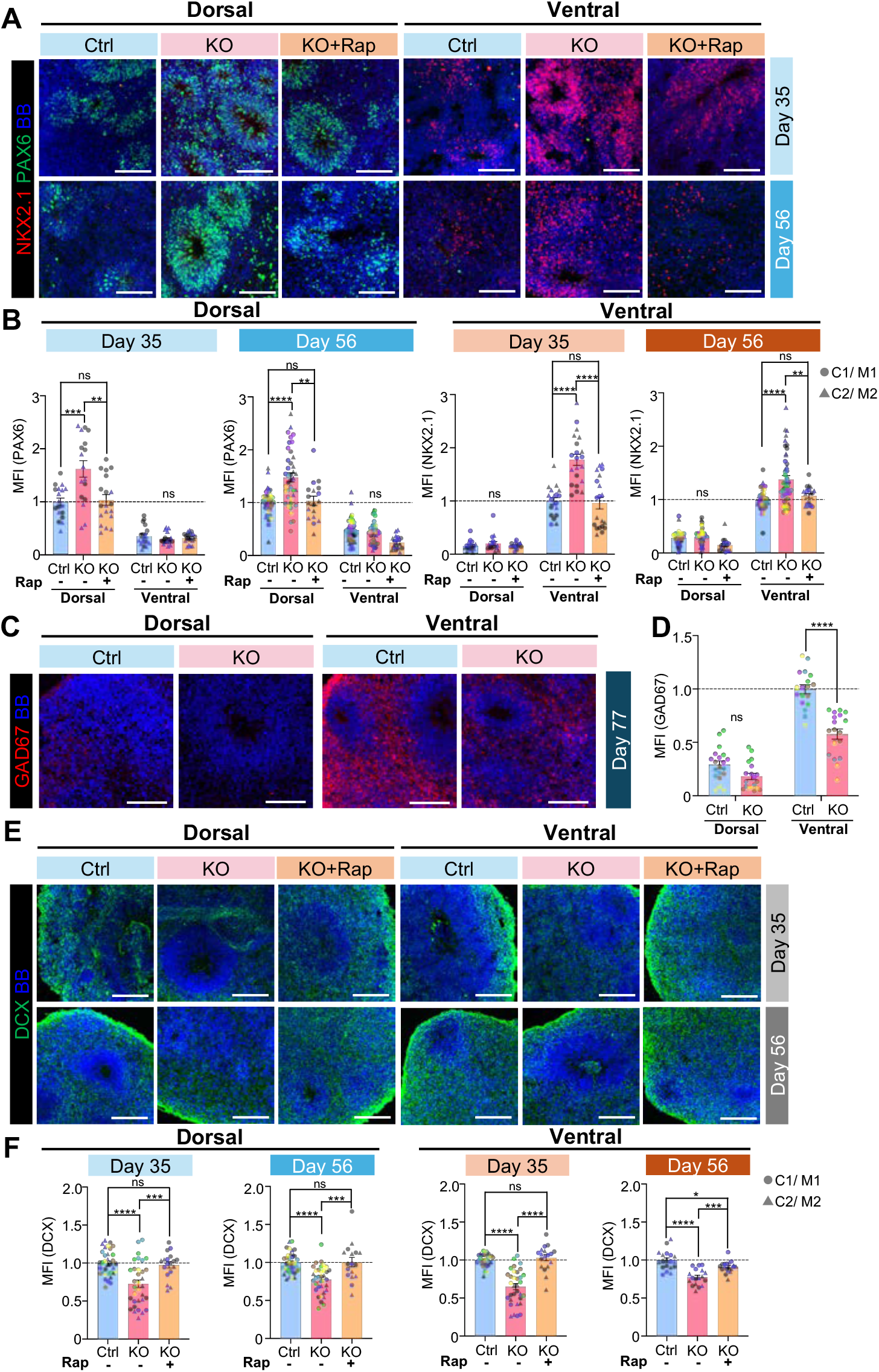
Forebrain regionalization in control and STRADA KO dhCOs and vhCOs. **(A)** Immunostaining for PAX6 and NKX2.1 in day 35 and 56 control and STRADA KO dhCOs and vhCOs treated with vehicle or rapamycin. **(B)** Quantification of relative PAX6 and NKX2.1 signals from A. **(C)** Immunostaining for GAD67 in day 77 control and STRADA KO dhCOs and vhCOs, with vehicle or rapamycin treatment. **(D)** Quantification of relative GAD67 signal for dhCOs and vhCOs from C. **(E)** Immunostaining for DCX in day 35 and 56 control and STRADA KO dhCOs and vhCOs treated with vehicle or rapamycin. **(F)** Quantification of relative DCX signal from E. Quantifications are computed as mean fluorescence intensity (MFI) within the organoid area. Each shape represents one organoid, compiled from separate batches of differentiation (color-coded). Mean ± SEM is displayed, with Mann-Whitney test performed for two-group comparison and one-way ANOVA for comparisons among three conditions. \**p*<0.05, \*\**p*<0.01, \*\*\**p*<0.001, \*\*\*\**p*<0.0001. Scale bars, A/C/E-100 µm.

Examination of the newborn neuron marker doublecortin (DCX) at days 35 and 56 revealed robust neurogenesis across all groups. There was a consistent decrease of DCX signal in KO dhCOs and vhCOs at both timepoints, which was rescued by rapamycin treatment (Fig. 3E–F). Nevertheless, DCX levels normalized between *STRADA* KO and control organoids by day 77 (data not shown), indicating that neurogenesis ultimately progresses despite early delays and similar to what we observed previously using a different dhCO protocol.^21^

### STRADA KO dhCOs exhibit delayed neurogenesis and increased neural progenitor populations

To better understand the effects of STRADA loss and mTOR hyperactivity on dorsal corticogenesis, dhCOs were collected at day 35, 56, and 77 to analyze cell fate differences during early neurogenesis, deep layer neuron production, and later stage generation of upper layer neurons, respectively.

Consistent with earlier findings (Fig. 3), STRADA KO dhCOs showed increased levels of the neural stem cell marker SOX2 and reduced expression of the mature neuron marker MAP2ab at day 35 compared to control (*p*<0.0001, Fig. 4A-B). With STRADA loss, the delay in neurogenesis, as shown by decreased MAP2ab signal, persisted at day 56 (Fig. 4A-B). Rapamycin treatment partially normalized SOX2 and MAP2ab expression (Fig. 4A-B).

**Figure 4.**
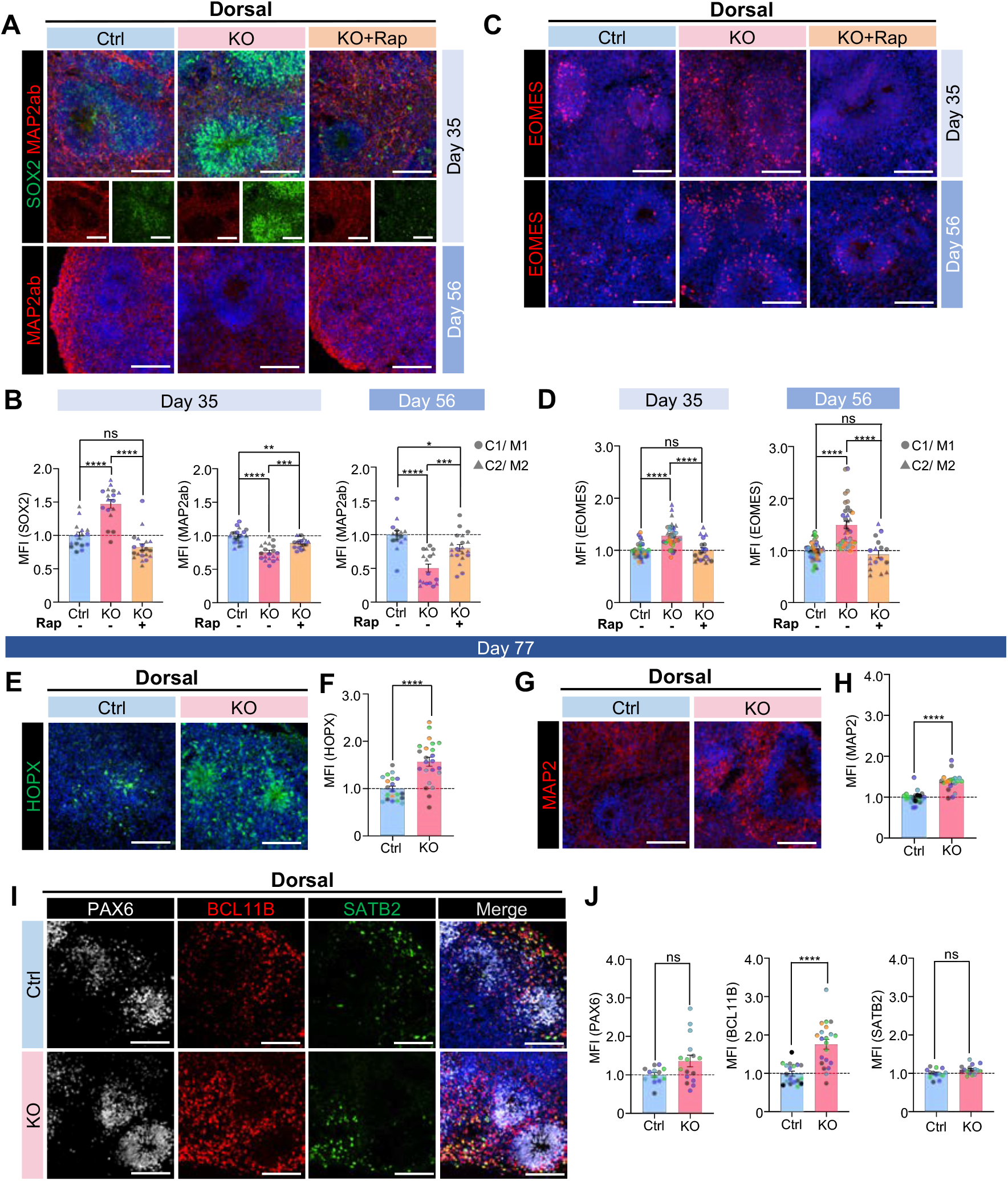
STRADA KO dhCOs show increased neural progenitors and delayed neurogenesis. **(A)** Immunostaining for SOX2 and/or MAP2ab in day 35 and 56 control and STRADA KO dhCOs treated with vehicle or chronic rapamycin. **(B)** Quantification of relative SOX2 and MAP2ab signals from A. **(C)** Immunostaining for EOMES in day 35 and 56 control and STRADA KO dhCOs treated with vehicle or chronic rapamycin. **(D)** Quantification of relative EOMES signal from C. **(E)** Immunostaining for HOPX in day 77 control and STRADA KO dhCOs. **(F)** Quantification of relative HOPX signal from E. **(G)** Immunostaining for MAP2 in day 77 control and STRADA KO dhCOs. **(H)** Quantification of relative MAP2 signal from G. **(I)** Immunostaining for PAX6, SATB2, and BCL11B in day 77 control and STRADA KO dhCOs. **(J)** Quantification of relative PAX6, SATB2, and BCL11B signals from I. Signal was measured as mean fluorescence intensity (MFI) within the area of each organoid. Each shape represents one organoid, compiled from separate batches of differentiation (color-coded). Data are shown as mean ± SEM. Statistical analysis was performed using the Mann-Whitney test for two-group comparisons and one-way ANOVA for comparisons among more than two conditions. \**p*<0.05, \*\**p*<0.01, \*\*\**p*<0.001, \*\*\*\**p*<0.0001. Scale bars, A/C/E/G/I-100 µm.

To assess whether the delay extended to intermediate progenitors (IPs), we stained for EOMES (TBR2), a marker of excitatory lineage IPs. STRADA KO dhCOs displayed significantly increased numbers of EOMES+ cells at both day 35 and 56 (Fig. 4C-D), and this was rescued by rapamycin treatment.

At day 77, we also examined HOPX+ oRGs and observed increased expression of HOPX in STRADA KO dhCOs at day 77 (4E-F), consistent with prior findings that mTORC1 hyperactivity promotes oRG fate.^21,29,30^ Alongside this, STRADA KO dhCOs exhibited higher overall levels of MAP2 (Fig. 4G-H), suggesting that the expanded progenitor pool ultimately results in greater neuronal output, correlating with the enlarged organoid size (Fig. 1B-C) and paralleling the megalencephaly phenotype seen in PMSE syndrome.

To determine whether ongoing changes in cell fate specification during neurogenesis at day 32 or only earlier changes in radial glia (RG) are contributing to the observed differences in cell markers in STRADA KO dhCOs, we labeled dividing cells with EdU at day 32 and analyzed cell populations at day 35 including the proportions of PAX6+/EdU+ (cycling ventricular RG), EOMES+/EdU+ (dividing cells that stay as or become IPs), and BCL11B+/EdU+ cells (dividing cells at day 32 that have undergone terminal differentiation into deep-layer neurons by day 35) (Fig. S4A-B). STRADA KO dhCOs displayed an enrichment of ventricular RG and IPs, with no difference in the number of newborn deep layer neurons, consistent with delayed neurogenesis and increased stem cell renewal in STRADA KO organoids.

Finally, we compared the distribution of PAX6+ neural progenitors, early born BCL11B+ deep layer neurons, and later born SATB2+ upper layer neurons in day 77 dhCOs to gauge the impact of delayed neurogenesis on terminal cell fate specification. By day 77, PAX6+ progenitor levels were comparable between control and KO dhCOs (Fig. 4I-J). We observed lower levels of earlier-born BCL11B+ deep layer neurons in KO dhCOs at day 56 (data not shown), consistent with delayed neurogenesis in KO organoids. However, by day 77 there was an increase in BCL11B+ deep layer neurons in KO dhCOs compared to control (Fig. 4I-J), suggesting that the increased neural progenitor pool in STRADA KO organoids eventually leads to increased output of deep layer neurons. There was no difference in later-born SATB2+ upper layer neurons in KO dhCOs compared to controls (Fig. 4I-J), despite the increase in HOPX+ progenitors in KO dhCOs (Fig. 4E-F).

### Ventral organoids exhibit delayed neurogenesis and increased progenitor populations in response to STRADA loss

We next analyzed control and STRADA KO vhCOs for altered neurogenesis in ventral forebrain development. vhCOs were collected at day 35, 56, and 77 to analyze cell fate differences during early, middle, and later stage interneuron development. STRADA KO vhCOs showed increased levels of the neural stem cell marker SOX2 and decreased levels of the mature neuronal marker MAP2ab at day 35 compared to control (Fig. 5A-B). The delay in neurogenesis as shown by decrease in MAP2ab signal in STRADA KO vhCOs was still apparent at day 56 (Fig. 5A-B). Rapamycin treatment in STRADA KO vhCOs normalized SOX2 and MAP2ab expression levels toward control values (Fig. 5A-B).

**Figure 5.**
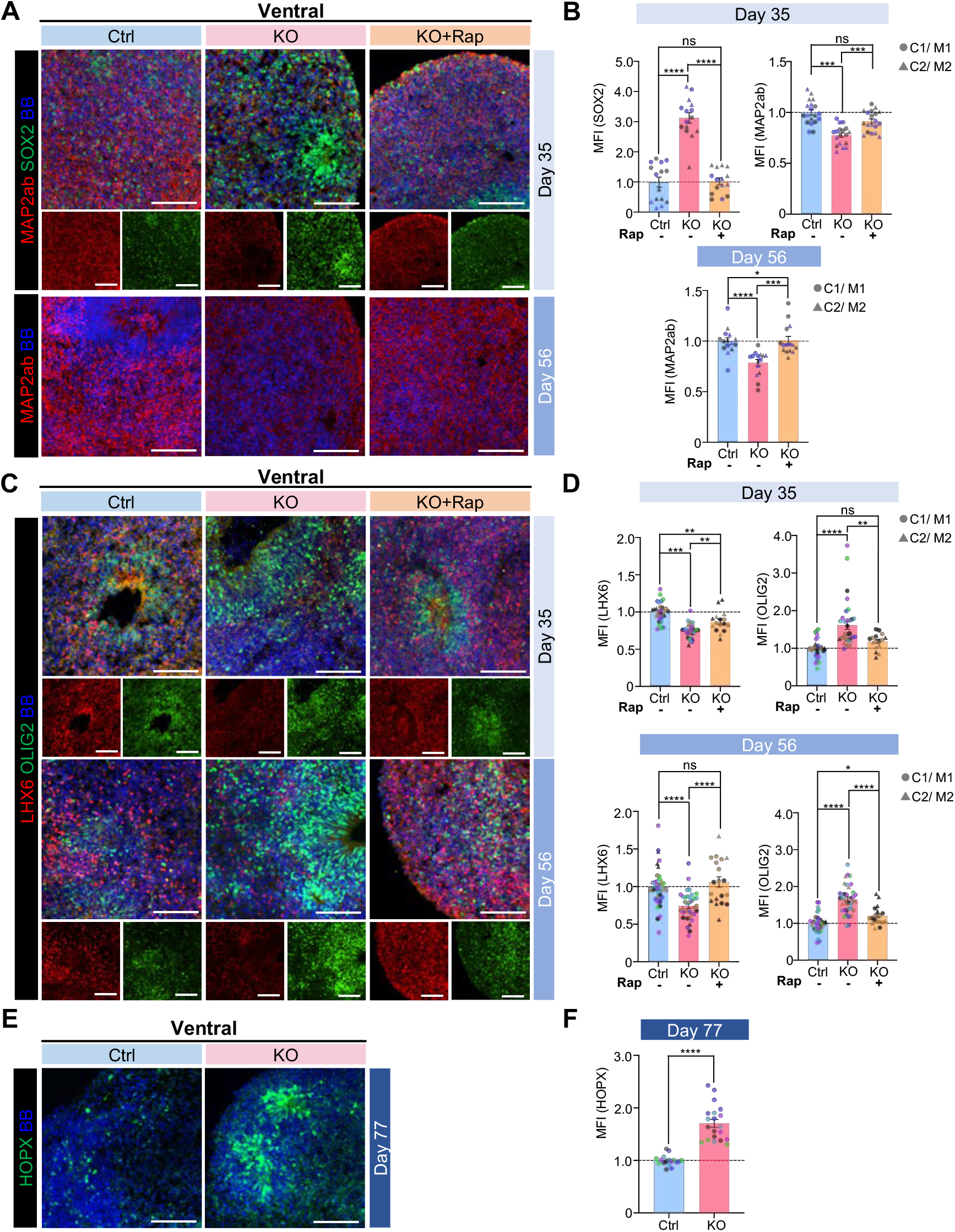
STRADA KO vhCOs exhibit increased neural progenitors and delayed neurogenesis. **(A)** Immunostaining for SOX2 and MAP2ab in day 35 and 56 control and STRADA KO vhCOs treated with vehicle or rapamycin. **(B)** Quantification of relative SOX2 and MAP2ab signals from A. **(C)** Immunostaining for OLIG2 and LHX6 in day 35 and 56 control and STRADA KO vhCOs treated with vehicle or rapamycin. **(D)** Quantification of relative OLIG2 and LHX6 signals from C. **(E)** Immunostaining for HOPX in day 77 control and STRADA KO vhCOs. **(F)** Quantification of relative HOPX signal from E. For all quantifications, signal was measured as mean fluorescence intensity (MFI) within the area of each organoid. Each shape represents one organoid, compiled from separate batches of differentiation (color-coded). Mean ± SEM is displayed, with Mann-Whitney test conducted for two-group comparison and one-way ANOVA for comparisons among more than two conditions. \**p*<0.05, \*\**p*<0.01, \*\*\**p*<0.001, \*\*\*\**p*<0.0001.

To further characterize ventral progenitor cell populations, vhCOs were stained for the early medial ganglionic eminence (MGE) progenitor marker OLIG2 and the MGE interneuron precursor marker LHX6. STRADA KO vhCOs showed significantly increased levels of OLIG2+ and significantly decreased levels of LHX6+ cells at both day 35 and 56 (Fig. 5C-D), consistent with a delay in progenitor maturation and neuronal differentiation. Treatment of KO vhCOs with rapamycin decreased the number of OLIG2+ progenitors and increased the number of LHX6+ neurons in STRADA KO organoids, suggesting that mTOR hyperactivity can cause delayed neurogenesis in ventral forebrain development (Fig. 5C-D).

To determine whether ongoing delayed neurogenesis at day 32 contributes to the observed differences in cell type composition in STRADA KO vhCOs, we performed EdU labeling at day 32 and quantified SOX2+/EdU+ progenitors and NeuN+/EdU+ neurons at day 35 (Fig. S1). STRADA KO vhCOs showed increased proportions of cycling SOX2+ progenitors compared to controls, a phenotype that was partially rescued by rapamycin (Fig. S4C-D). The number of newly generated NeuN+ neurons, however, remained comparable between groups at this early stage (Fig. S4C-D), indicating that increased progenitor retention rather than complete neurogenesis arrest characterizes the early phenotype.

We also examined the levels of HOPX+ oRGs in control and KO vhCOs at day 77, as HOPX+ progenitors are known to be present in the ganglionic eminences.^31^ We observed increased HOPX+ progenitors in KO vhCOs (Fig. 5E-F), suggesting that the bias toward progenitor renewal extends into the later stage of ventral neurogenesis. Taken together, these results demonstrate that loss of STRADA disrupts ventral forebrain development via prolonged retention of ventral progenitors and delayed interneuron differentiation in an mTOR-dependent manner, consistent with the broader disruption of neurogenesis observed across forebrain regions.

### STRADA KO vhCOs showed altered interneuron subtype specification

To assess the effects of STRADA loss and mTORC1 hyperactivity on interneuron subtype specification, we analyzed vhCOs at days 56 and 77 for markers of inhibitory neuron subtypes. Immunostaining was performed for GABA, a broad marker of GABAergic inhibitory interneurons, as well as for interneuron subtype markers including calretinin (CALRET), calbindin (CALB), somatostatin (SST), and neuropeptide Y (NPY).^32–35^

Calretinin-positive interneuron numbers were similar between control and STRADA KO vhCOs at both day 56 and day 77 (Fig. 6A, 6I-J). However, STRADA KO vhCOs showed a significant decrease in calbindin- and SST-positive interneurons relative to controls at both day 56 and 77 (Fig. 6C–F, 6I-J). Rapamycin treatment partially rescued the deficit in SST+ interneurons at day 56 but had minimal effect on calbindin levels (Fig. 6I). Notably, the numbers of NPY-positive interneurons were comparable between control and KO at day 56 but exhibited a modest increase in KO vhCOs by day 77 (Fig. 6I-J). Additionally, consistent with their prolonged maturation timeline, parvalbumin (PV)-positive interneurons were not observed at day 56 or 77. Future studies extending to later developmental stages will be important to fully capture the emergence of PV-positive interneurons and to assess how prolonged mTOR hyperactivity impacts their maturation and function.

**Figure 6.**
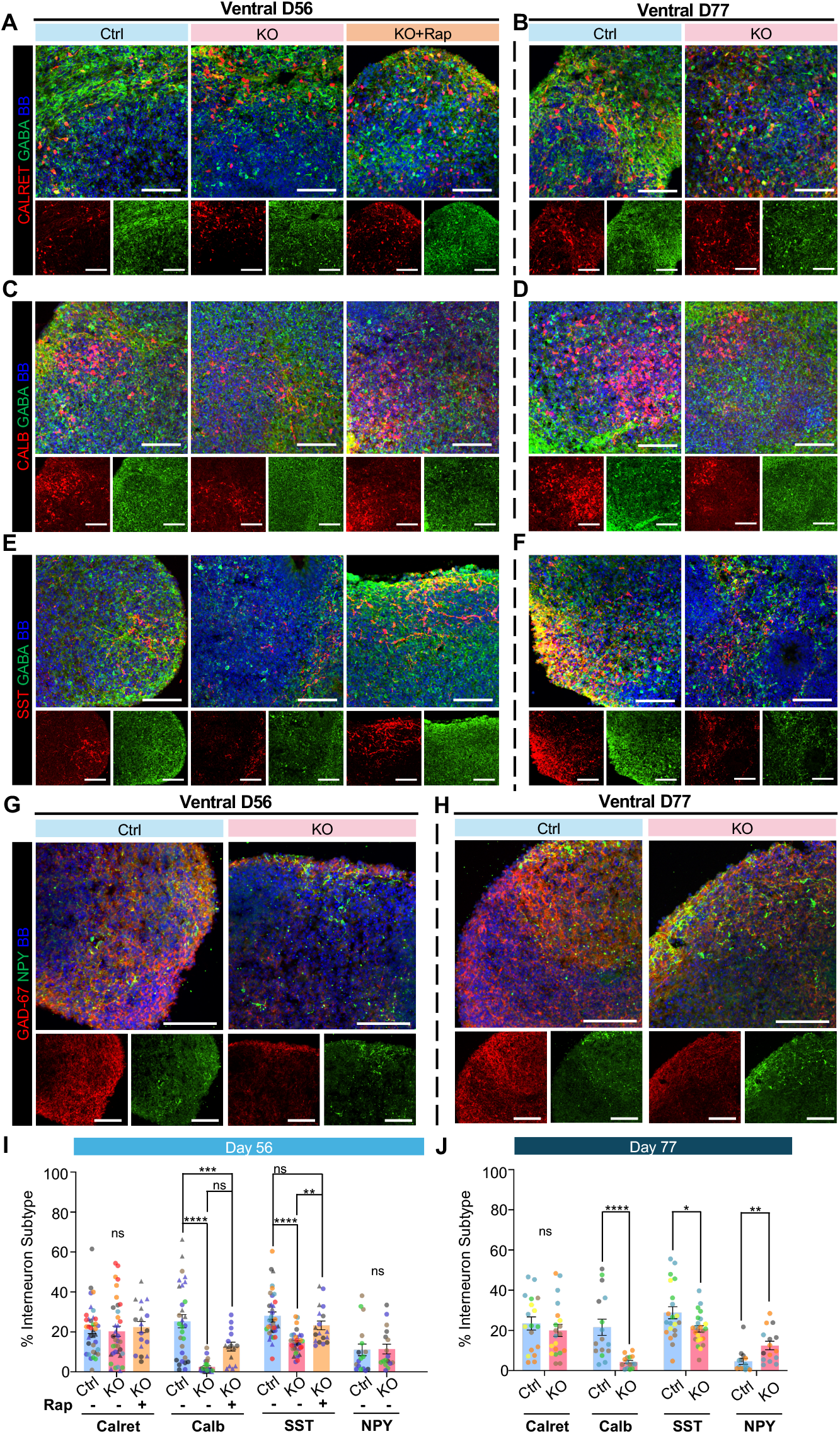
STRADA loss drives altered interneuron specification. **(A-H)** Representative immunostaining of day 56 (A, C, E, G) and day 77 (B, D, F, H) vhCOs for multiple subtypes of GABAergic inhibitory interneurons: Calretinin (A, B), calbindin (C, D), somatostatin (E, F), and neuropeptide-Y (G, H). The first three were co-stained with GABA, and neuropeptide-Y was co-stained with GAD-67. Treatment with rapamycin was included for day 56 organoids. Scale bars, 100 µm. **(I-J)** Quantification of interneuron subtypes from day 56 (I) and day 77 (J) vhCOs, expressed as the proportion of GABAergic cells positive for each interneuron marker. Each symbol represents one organoid from independent differentiation batches (color-coded). Data are presented as mean ± SEM; statistical comparisons were performed using either Mann-Whitney test) (J) or one-way ANOVA (I). \**p*<0.05, \*\**p*<0.01, \*\*\**p*<0.001, \*\*\*\**p*<0.0001.

Single cell RNA-sequencing analysis reveals altered cell fate specification and differentially expressed genes in dorsal and ventral STRADA KO organoids

We used single cell RNA-sequencing (scRNA-seq) to analyze the cell-type specific genome-wide transcriptomic effects of STRADA loss of function on dorsal and ventral forebrain development and assess for shifts in cell type composition that can result from altered cell fate. Samples for scRNA-seq are listed in Table 1. Data from dhCOs and vhCOs were separately analyzed by unsupervised clustering in Uniform Manifold Approximation and Projection (UMAP) plots, combining data from day 35 and day 77 (Fig. 7). For dhCOs, unsupervised clustering defined 24 clusters, and after additional filtering, manual curation defined 12 clusters (Fig. S5A-B, 7A). Cortical RG were marked by *PAX6, SOX2, VIM*, and *FABP7*, dividing cells by *MKI67* and *NUSAP1*, IPs by *EOMES*, oRG by *HOPX*, and excitatory neurons by *STMN2, GAB43, SLC17A7* (VGLUT1) and *SLC17A6* (VGLUT2) (Fig. S5C-D). Metabolically active RG were distinguished by higher expression of genes encoding ribosomal proteins. There were also substantial clusters of ganglionic eminence (GE) marked by *DLX1, DLX2, DLX5*, and *DLX6*, and midbrain cells marked by *NTRK2, PAX7, LHX1* and *LMX1B* expression and *FOXG1* negativity. GE and midbrain cells were mostly present in the day 35 compared to the day 77 samples (data not shown).

**Figure 7.**
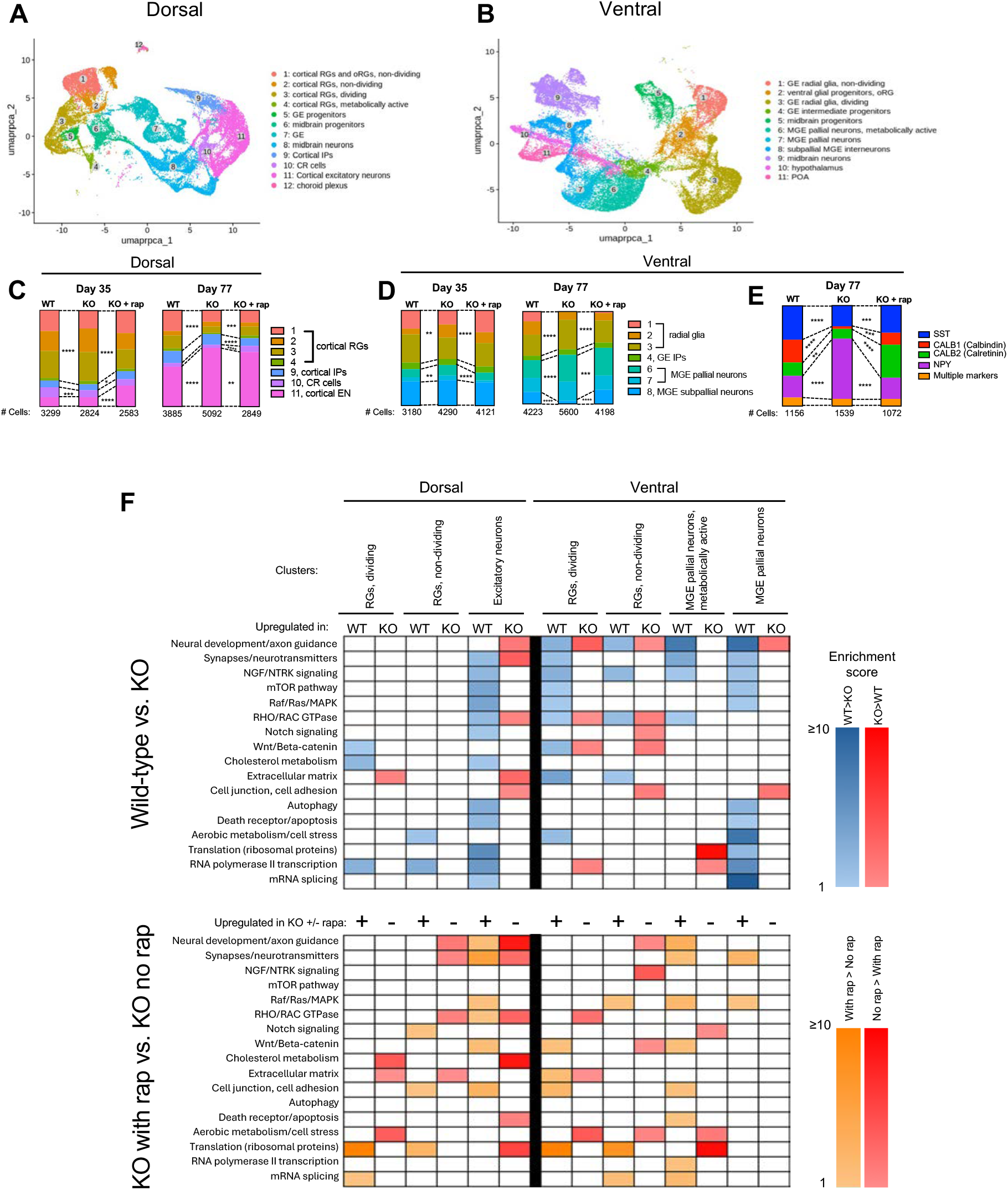
scRNA-Seq shows altered neurogenesis and cell fate specification in STRADA KO dhCOs and vhCOs compared to control. **(A-B)** Combined UMAP plots of all dhCO (A) and vhCO (B) samples with cluster identification. Legend at right displays the cell type identity for each cluster. **(C)** Bar graphs showing the percentage of cells found in selected clusters of control and STRADA KO dhCOs. **(D)** Bar graphs showing the percentage of cells found in selected cluster of control and STRADA KO vhCOs. **(E)** Differentially represented clusters of pathways between WT and KO and between KO with rapamycin and KO without rapamycin from selected cell type clusters.

The proportions of cortical cells (RGs, IPs, Cajal Retzius cells, and excitatory neurons) in dhCOs was compared in each condition, at day 35 and day 77. At day 35 when more progenitors are present, there were increased cortical RGs in KO compared to WT, and this was rescued with rapamycin (Fig. 7C). Conversely, at day 77, there were increased excitatory neurons and decreased RG in KO compared to WT, and this was also rescued by rapamycin. This is consistent with a retained stem cell fate and expanded progenitor pool in early cortical development, resulting in an overproduction of neurons later in development.

For vhCOs, unsupervised clustering defined 18 clusters, and filtering followed by manual curation identified 11 clusters (Fig. S6A-B, 7B). Ventral forebrain RG were marked by *SOX2, VIM, NKX2-1* positivity and *PAX6* negativity. Dividing cells were marked by *MKI67* and *NUSAP1*. Ventral intermediate progenitors were marked *by DLL1, DLL3* and *ASCL1*. MGE markers *NKX2-1, DLX1, DLX2, DLX5, DLX6*, and *LHX6* were present, while caudal GE (CGE) markers including *NR2F2* and *SCGN* were mostly absent (Fig. S6C-D). Subpallial/striatal MGE neurons were marked by *LHX8*, while pallial MGE neurons were marked by *LHX6*. MGE-derived pallial interneurons were divided into two clusters (cluster 6 and 7 in Fig. S6B), distinguished by higher expression of genes encoding ribosomal proteins in the “metabolically active” cluster 6. Inhibitory neurons were marked by *GAD1* and *SLC32A1*, and there were only a small population of *SLC17A6* excitatory neurons in the hypothalamus cluster. Preoptic area was marked by *HMX3*, hypothalamus by multiple markers including *ISL1* and *GNRH*. Midbrain cells were marked by *NTRK2, NKX6-1, LHX1*, and *LMX1B* expression, and lack of *FOXG1*.

The proportion of GE cells in vhCOs was compared in day 35 and day 77 (Fig. 7D). At day 35, there was a slight decrease in the proportion of RG and a slight increase in MGE pallial neurons in KO compared to WT control. These changes were rescued by rapamycin. At day 77, there was also a decrease in RG, increase in MGE pallial neurons, and decrease in MGE subpallial neurons in KO compared to WT control. With rapamycin treatment of day 77 KO organoids, the proportion of RG decreased further, and the amount of GE pallial neurons was slightly lower as well, with an increase in MGE subpallial neurons.

Proportions of interneuron subtypes in vhCOs at day 77 were determined by examining known interneuron markers *SST, CALB1, CALB2, NPY, CCK, PVALB, HTR3A*, and *VIP*. Very few cells expressed *CCK, VIP, HTR3A*, or *PVALB* (<2%). *SST, CALB1, CALB2* and *NPY* were expressed in vhCOs (and 7-9% of cells expressed multiple of these markers), with KO vhCOs having fewer *SST, CALB1* (Calbindin), and *CALB2* (Calretinin) neurons, and increased *NPY+* neurons compared to WT, and with rapamycin rescue of these changes (Fig. 7E).

Next, we determined whether STRADA WT vs. KO and rapamycin treated vs. vehicle treated KO hCOs had altered transcriptomic signatures by analyzing cell-type specific differentially expressed genes (DEGs). For each cell type and comparison, DEGs were entered into the Database for Annotation, Visualization, and Integrated Discovery (DAVID) Bioinformatics tool^36,37^ to find overrepresented Reactome pathways, and clustering of the pathways was performed on select cell types to group pathways that shared similar sets of DEGs. These clustered pathways were categorized representatively, and categories of pathways that were noted repeatedly in the comparisons included aerobic metabolism, protein translation, mRNA transcription and splicing, cell signaling (NTRK, mTOR, Raf/Ras/MAPK, Rho/Rac GTPase, Notch, Wnt/Beta-catenin), cholesterol metabolism, neuronal development, autophagy, and cell death (Fig. 7F). In some instances, upregulated genes in contrasting conditions were components of the same pathways, therefore those pathways were overrepresented in both conditions (e.g. ventral dividing RG, WT vs. KO and neural development/axon guidance).

## DISCUSSION

### STRADA KO vhCOs provide novel insight into roles of interneuron development in mTORopathy pathogenesis

While mTORopathies are a significant cause of cortical malformations and neurological comorbidities, their genetic and phenotypic heterogeneity makes the study complicated. Focal cortical dysplasia (FCD), often requiring resective epilepsy surgery, provides access to dorsal forebrain progenitor-derived tissue carrying mTOR mutations. In our case, all brain cells carry the pathogenic variant, but the lack of surgical indication for megalencephaly limits study of ventral forebrain-derived interneurons. Although altered interneuron populations have been reported in some mTORopathies like FCD Type II^38–42^, most human tissue comes from post-developmental stages after years of chronic seizures. Animal models offer mechanistic insights^27,43^, but differences in brain development,^29^ interneuron diversity,^44^ and somatic mosaicism^45,46^ have limited their translational relevance to the corresponding human disorders. In all, our model addresses these gaps by enabling analysis of how mTORC1 hyperactivity affects interneuron development in a genetically relevant context.

More recent undirected cerebral organoid models of mTORopathies like TSC have implicated CGE-lineage interneurons in early lesion development, with later involvement of dysmorphic excitatory neurons and minimal MGE contribution.^47^ However, given that a substantial portion of cortical interneurons in humans originate from the MGE, and that chronic mTORC1 hyperactivity—as observed in disorders like PMSE—affects all progenitor domains, it is likely that MGE development and its derived interneurons also contribute significantly to pathogenesis. Our work broadens this perspective by examining interneurons from various lineages and therefore reveals additional mechanisms by which mTOR dysregulation disrupts subpallial development.

Taken together, our study establishes a human-derived platform enabling direct investigation of aberrant early neurodevelopment in response to STRADA loss and mTOR hyperactivity, which cannot be fully recapitulated in animal models. By incorporating region-specific patterning of ventral forebrain, we reveal previously underrecognized roles of MGE-derived interneurons in the pathogenesis of PMSE, raising the possibility that widespread aberrant interneuron development contributes to the significantly more severe epilepsy and cognitive phenotypes seen in PMSE compared to FCD or TSC.

### Hyperactive mTORC1 signaling results in increased progenitor pools and delayed neuronal differentiation in both dhCOs and vhCOs

A consistent finding across our model was that STRADA loss led to increased maintenance of neural progenitor (neural stem cell and intermediate progenitor) fate and delayed neuronal differentiation in both dhCOs and vhCOs. This expanded progenitor pool during early development in STRADA KO resulted in an increase in neuronal numbers later in development, recapitulating megalencephaly seen in PMSE.

While it remains unclear how closely the delayed neurogenesis observed in our mutant organoids mirrors *in vivo* developmental timing in PMSE, even subtle disruptions during early differentiation may have cascading effects on subsequent development. These could include alterations in the balance and identity of mature neuronal subtypes and impairments in subsequent neuronal migration^48,49^ and connectivity,^50^ ultimately contributing to the widespread network dysfunction seen in PMSE patients.

It remains unknown whether dorsal and ventral organoids are equally affected by mTORC1 hyperactivation or whether the delay occurs with differing magnitudes or kinetics between regions. Notably, vhCOs typically exhibit a more prolonged developmental timeline compared to dhCOs, to generate functionally mature interneurons.^51,52^ As such, mTORC1-driven delays may further desynchronize the developmental progression between dorsal and ventral forebrains, potentially exacerbating defects in excitatory-inhibitory integration and circuit formation.

We also observed increased cell proliferation in STRADA KO organoids, consistent with known effects of mTORC1 hyperactivity. Interestingly, the early decrease in cell death at day 14 was not rescued by chronic rapamycin treatment, suggesting a more complex role for STRADA signaling in regulating apoptosis during brain development. This aligns with prior findings that apoptotic dysregulation in mTORopathies may involve mTORC1-independent mechanisms.^53^

Together, our findings suggest that delayed neurogenesis and prolonged progenitor fate may contribute not only to altered neuronal subtype composition, but also to broader disruptions in forebrain wiring and function. Future studies aiming at determining how mTORC1 signaling affects later developmental events, such as interneuron maturation and cortical integration, will be essential to understanding PMSE pathogenesis and identifying opportunities for targeted intervention.

### Hyperactive mTORC1 signaling alters cell fate specification of interneurons in STRADA KO vhCOs

Prior mouse studies have shown that mTOR signaling is essential for interneuron development. Early deletion of genes like *Pik3ca, Pten*, or *Tsc1* leads to mTORC1 hyperactivation and reduced interneuron density, mirroring findings in human mTORopathy tissues.^54–56^ Additionally, shifts in interneuron subtypes have been documented across various models: *Pten* deletion via Nkx2.1-Cre or Dlx1/2-Cre increases the PV:SST ratio,^55^ while Lhx6-Cre-mediated deletion increases both PV+ and SST+ subpopulations.^57^*Tsc1* deletion in SST+ cells resulted in PV expression and fast-spiking properties characteristic of PV cells.^58^

In our STRADA KO vhCO model, chronic mTORC1 hyperactivity led to a reduced number of SST+ and calbindin+ and increased NPY+ interneurons compared to isogenic controls. We did not detect parvalbumin (PV)+ interneurons at day 77, likely due to their later emergence in development. Therefore, it remains unclear whether a compensatory increase in PV+ cells occurs at later stages, as observed in some animal models.^55,57,59^ Future work will investigate PV+ interneuron maturation and overall subtype composition at extended timepoints.

Our finding of increased NPY+ interneurons in STRADA KO contrasts with reduced cortical NPY+ interneurons seen in *Tsc1* conditional KO mice using Dlx5/6-Cre.^56^ NPY+ interneurons are known to be morphologically, physiologically, and transcriptionally diverse^60^ and likely arise from distinct developmental origins across subtypes.^61,62^ While the expanded NPY+ population in our STRADA KO vhCOs is presumably MGE-derived, their precise lineage, maturation state, and electrophysiological properties remain to be defined. Given NPY’s roles in energy balance, stress response, memory, and endogenous anticonvulsant activity,^63,64^ persistent elevation of NPY+ interneurons in STRADA-deficient brain tissue could represent a compensatory or pathogenic mechanism in PMSE, again emphasizing the complex and context-specific effects of mTORC1 dysregulation on interneuron specification.

Overall, the functional consequences of altered interneuron subtype composition in PMSE remain unclear. However, disruptions in the balance of inhibitory subtypes—alongside defects in excitatory neuron development—likely contribute to dysfunctional synaptic connectivity and miswiring of cortical circuits. Furthermore, delayed maturation of specific interneuron populations, or inhibitory neurons broadly, may interfere with proper network formation. Such mismatches in the timing of excitatory-inhibitory development could underlie the neurodevelopmental features of PMSE, including epilepsy and cognitive impairment.

### scRNA-seq analysis provides insight into potential mechanisms of PMSE pathogenesis

At day 35, cell type composition of the dhCOs but not the vhCOs by scRNA-seq matched data from immunohistochemistry (IHC) analysis that showed delayed neurogenesis in KO vs. WT. The difference in the findings in vhCOs may be explained by the difference in number of replicates (2-6 differentiations in IHC compared to 2-3 differentiations in scRNAseq) and the differences between cell type assignment by transcriptomics compared to measuring protein levels of known cell type markers in IHC. At day 77, both dhCOs and vhCOs had increased percentage of neurons comprising the organoids in KO vs. WT, consistent with prior expansion of the progenitor pool from earlier delayed neurogenesis, and this elucidates a potential mechanism for megalencephaly in PMSE.

The cell-type specific differentially represented clusters of pathways for KO vs. WT and KO without rapamycin vs. KO with rapamycin suggests underlying molecular mechanisms for pathogenesis in PMSE and mTOR-related megalencephaly. Altered Rho GTPase signaling and axon guidance pathways may be associated with deleterious effects on tangential migration of interneurons. Defective RhoA signaling and altered axon guidance has been demonstrated in TSC patient-derived iPSC neurons,^65^ and prior studies on STRADA-depleted mouse neural progenitor cells and PMSE patient-derived fibroblasts have demonstrated impaired migration, linked to disrupted Rho GTPase signaling and actin organization, downstream of mTORC1.^48^ Studying whether mTORC1 hyperactivity in STRADA results in impaired interneuron migration is a potential future direction of study. Gene expression in various signaling pathways known to affect neural stem cell self-renewal including Wnt/beta-catenin and Notch were altered, suggesting that the effects of mTORC1 hyperactivity may be mediated by other canonical developmental pathways or have crosstalk with other pathways in PMSE. Differentially represented synapse/neurotransmitter pathways suggest that for PMSE, aberrant mTORC1-mediated regulation of synaptogenesis may also be a mechanism for neuronal dysfunction that could result in epilepsy and cognitive impairment. Alterations in several metabolic and stress-related pathways including autophagy, apoptosis, cell stress, translation, and transcription are consistent with known functions of the mTORC1 and raise questions of how the downstream effects of mTORC1 hyperactivity result in altered brain development.

### Limitations of the study

Although multiple CRISPR-edited iPSC lines reduce background variability, it remains unclear whether similar phenotypes occur in organoids differentiated from PMSE patient-derived iPSCs. Limited differentiation replicates, especially for some treatment groups, constrain the generalizability of transcriptomic results. We did not include other pharmacological or genetic rescue strategies beyond rapamycin, which limits our ability to assess the robustness and specificity of the phenotypes. Importantly, there are likely mTOR-independent effects of STRADA loss through substrates of LKB1 and AMPK. Finally, inherent limitations of current cortical organoid models—including lack of full brain architecture, vasculature, and signaling gradients—warrant future use of advanced systems with better neuronal maturation (e.g., assembloids, patterned gradient culture systems) and comparative analyses with human tissue datasets.

### Data and code availability

- Single-cell RNA-seq data have been deposited at GEO at NCBI with accession number GSE296775.

## ACKNOWLEDGMENTS

The authors would like to thank Preethi Swaminathan for technical support and Jack Parent and the Parent laboratory for resources and insightful discussions on this project. We thank Whitney Parker and Wei Niu for their comments on the manuscript.

Library prep and next-generation sequencing was carried out in the Advanced Genomics Core at the University of Michigan. Research reported in this publication was supported by the University of Michigan Advanced Genomics Core, and the National Institutes of Health under Award Number P30CA046592 using the Cancer Center Single Cell and Spatial Analysis Shared Resource.

pX330-U6-Chimeric_BB-CBh-hSpCas9 was a gift from Feng Zhang.

This research was supported by the National Institutes of Health (grant number R01NS127829), and the Pediatrics Department at the University of Michigan, including the Charles Woodson Research Fund.

## AUTHOR CONTRIBUTIONS

Conceptualization, L.T.D., K.I., W.C., T.P., D.C.J., and G.L.; methodology, L.T.D., T.P., G.L., J.S., D.C.J., X.L., and W.C.; Investigation, T.P., G.L., X.L., D.V., J.C.W., S.K., A.S., A.K., D.C.J., S.P-G., S.V., J.S.; writing—original draft, G.L., T.P., D.V., J.C.W., X.L., W.C., and L.T.D.; writing—review & editing, G.L., T.P., L.T.D., X.L., K.I., and W.C.; funding acquisition, L.T.D.; supervision, L.T.D., W.C., and T.I.

## METHODS

### KEY RESOURCES

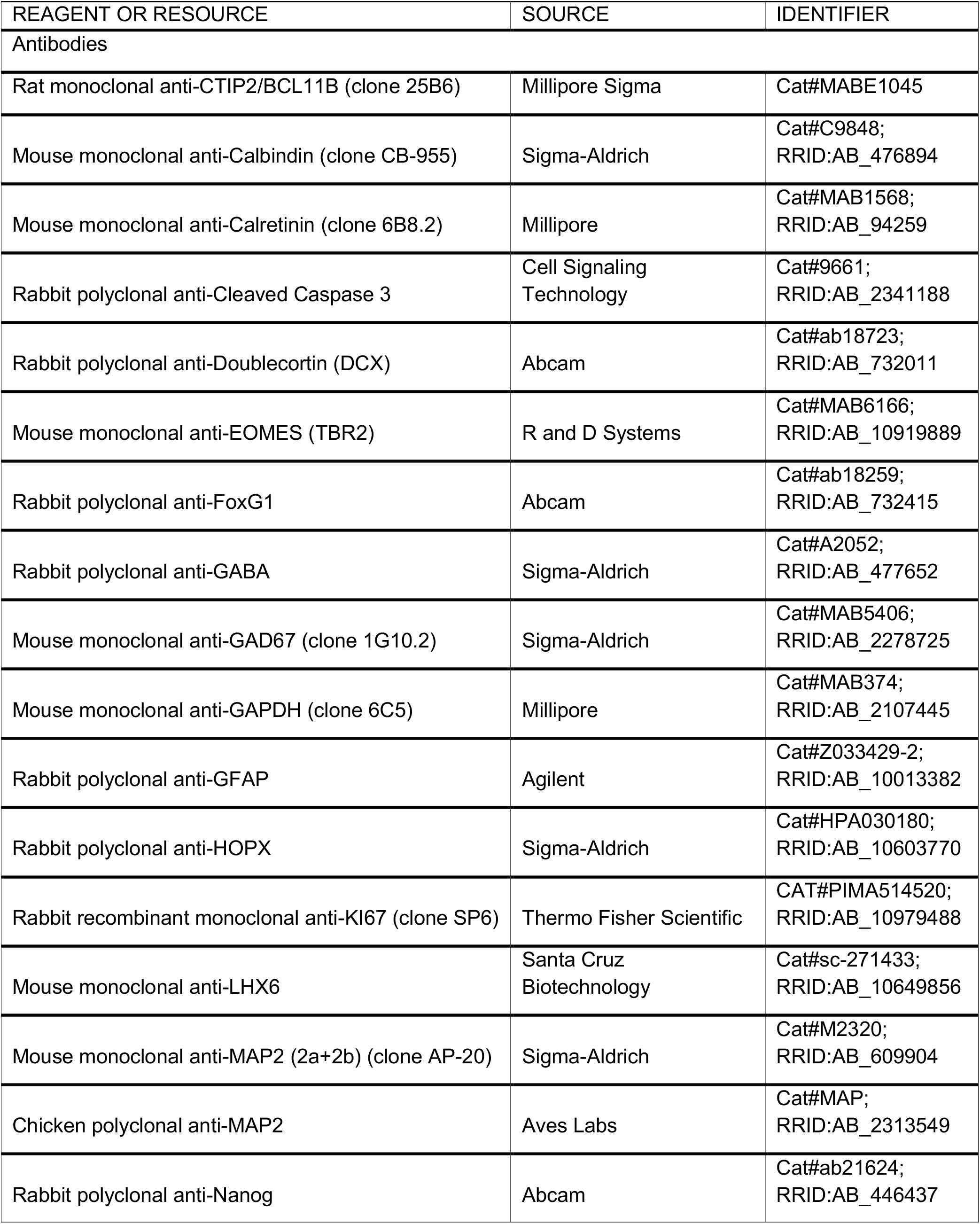
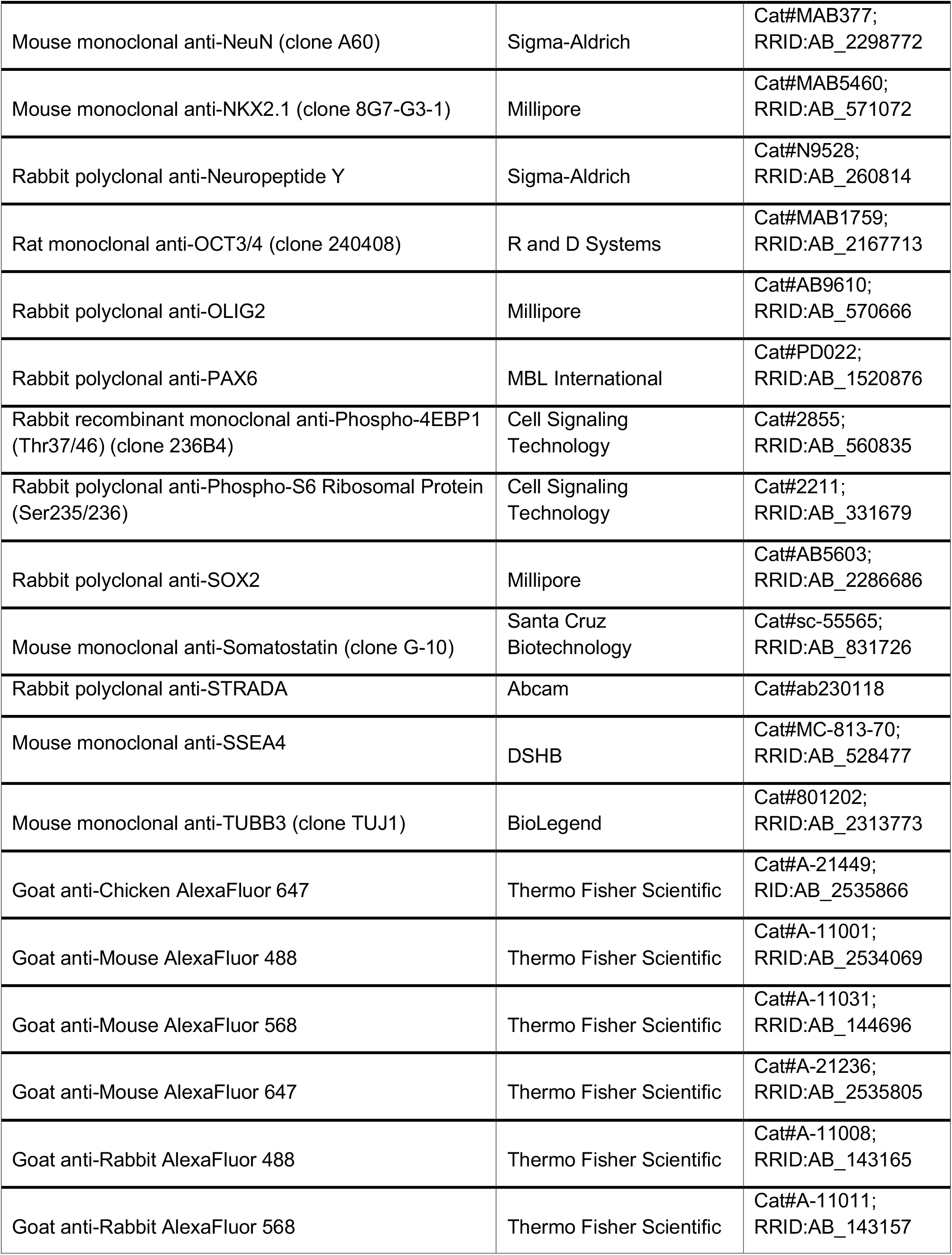
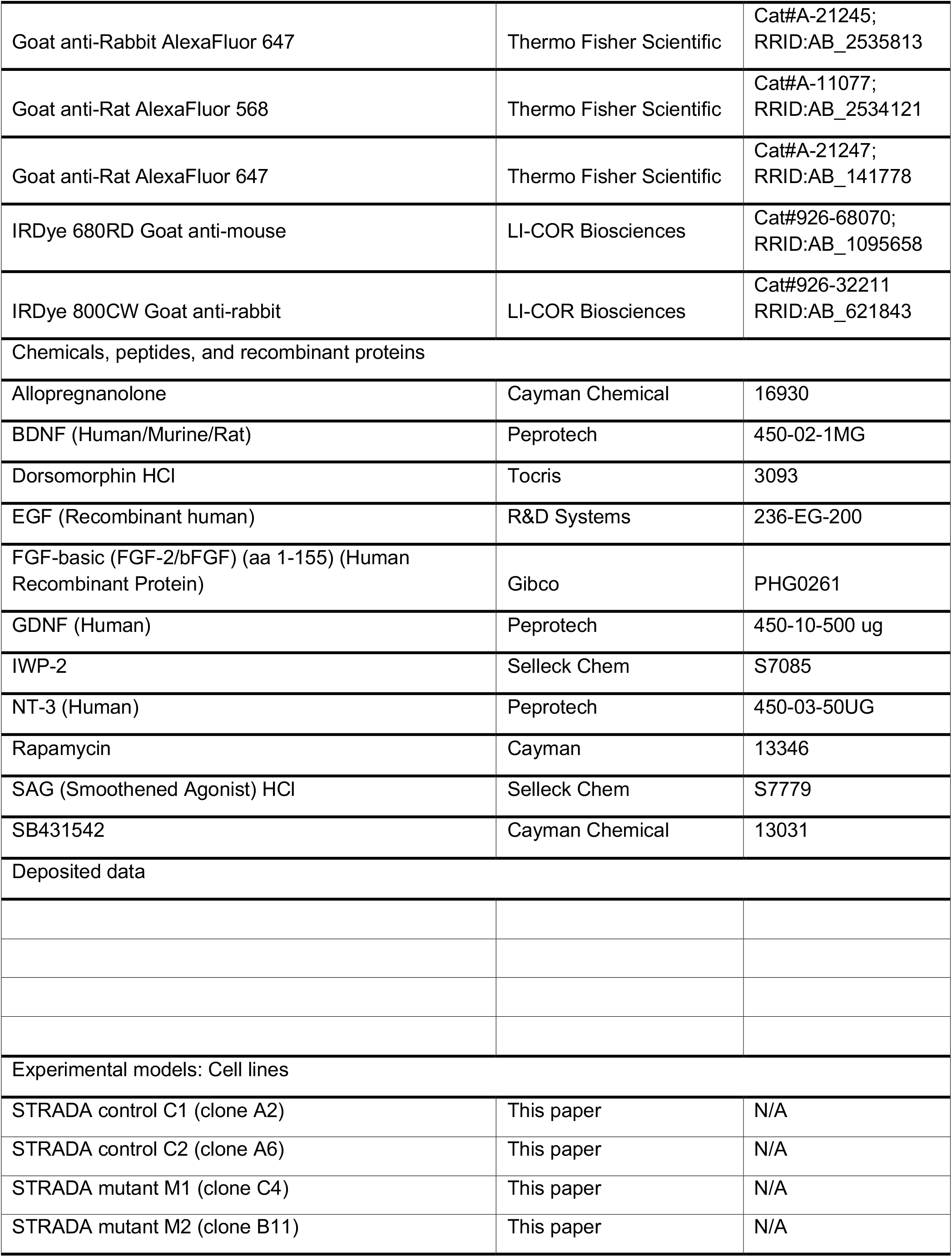
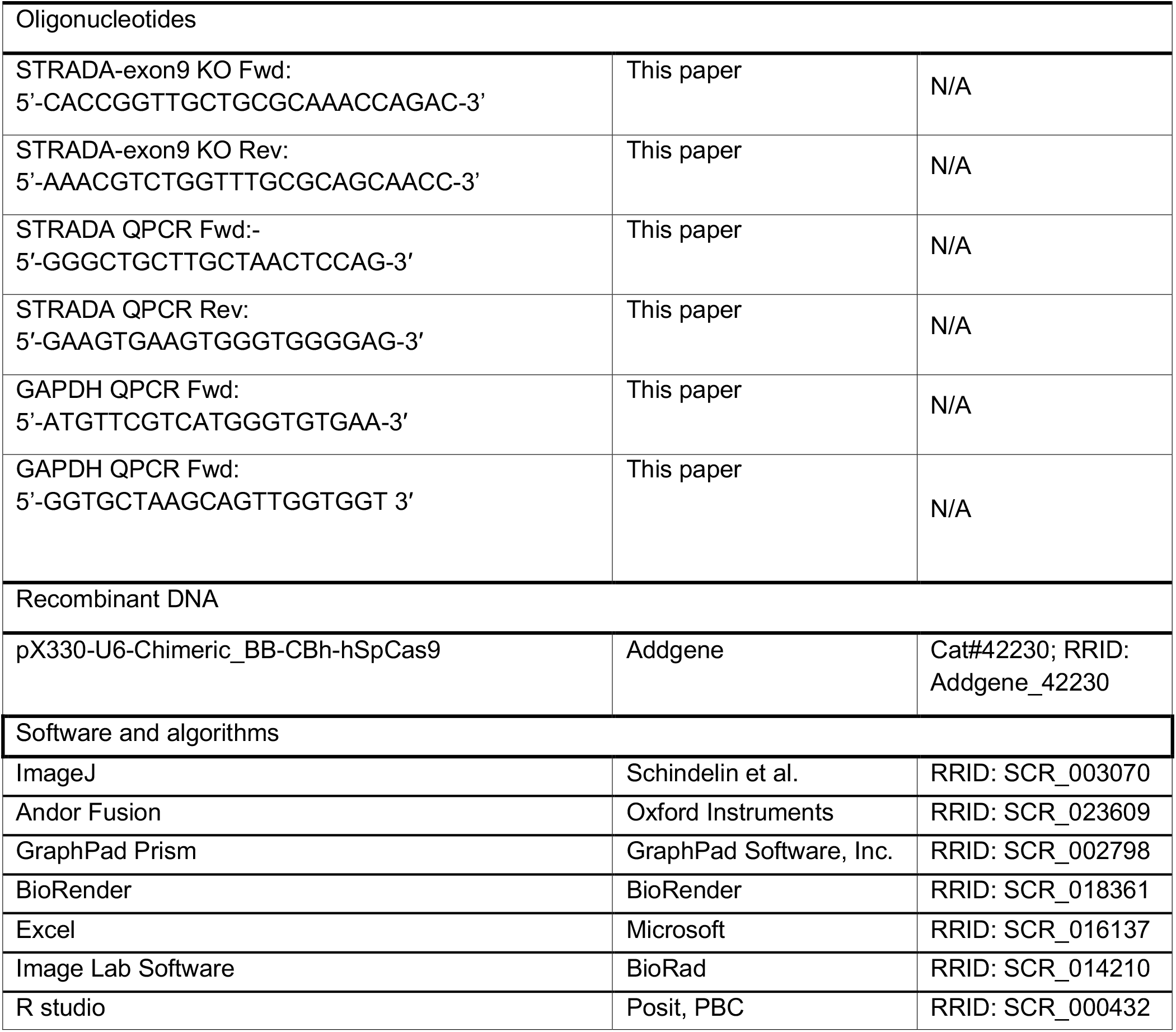

### EXPERIMENTAL MODEL AND STUDY PARTICIPANT DETAILS

The four iPSC lines used for this study (C1, C2, M1, M2) were derived from a commercially available source of male foreskin fibroblasts using CRISPR gene editing of *STRADA*. C1 and C2 lines were unedited and carried the wild-type sequence of STRADA (NM_001003787.4). M1 was a compound heterozygote, with c.632_636del (p.Leu211Argfs*45) on one allele and c.636del (p.Gly213Valfs*8) on the other. M2 was also a compound heterozygote, with two variants, c.635_636del (p.Ser212Trpfs*45) and c.631C>G (p.Leu211Val) on one allele, and c.636_637insA (p.Gly213Argfs*45) on the other allele.

### METHOD DETAILS

#### Cell line generation/Establishment of CRISPR KO Cell Lines

Oligonucleotides “STRADA-exon9 KO Fwd” and “STRADA-exon9 KO Rev” were annealed and cloned into pX330 (Addgene 42230) in generate gRNA targeting exon 9 of STRADA with CRISPR/Cas9.^67^ Foreskin fibroblast cells (NuFF, GlobalStem, Inc., GSC-3002) were simultaneously reprogrammed and CRISPR genome edited to generate STRADA knockout iPSC lines using a previously published protocol.^68^ Two *STRADA* WT clones were used as isogenic control lines (C1, C2) and two *STRADA* KO lines (M1, compound heterozygous mutations; M2, compound heterozygous mutations) from the same reprogramming experiment were used for subsequent experiments.

#### Induced pluripotent stem cell culture/iPSC line verification

Stem cell colonies were maintained at 37°C, 5% CO2, in mTeSR Plus media (Stemcell Technologies, Vancouver, British Columbia, Canada), on tissue culture plates pre-coated with Geltrex (1:200 dilution in DMEM/F12, Thermo Fisher, Waltham, Massachusetts, USA). iPSCs were passaged every 4–7 days using Dispase (Thermo Fisher Scientific, Waltham, Massachusetts, USA). Cultures were confirmed mycoplasma negative via periodic testing with a PCR-based assay. Each line used for experiments had expression of pluripotency markers (NANOG, OCT3/4, SSEA, SOX2). Reprogrammed lines tested negative for episomal reprogramming vector integration. Each line was assessed for chromosomal abnormalities by G-band karyotyping (Cell Line Genetics, Madison, Wisconsin, USA), single nucleotide polymorphism (SNP) chip microarray analysis (Infinium Core Exome-24 BeadChip, Illumina, San Diego, California, USA), or both.

#### Generation of cortical organoids

Cortical organoids were generated using a previously described protocol (Birey, Pasca 2017; Sloan, Pasca 2018; Miura, Pasca 2022) with minor modifications.^28,69,70^ On day -2, iPSCs were dissociated to single cells using Accutase (Innovative Cell Technologies, San Diego, California, USA), and plated in mTeSR Plus with 50 μM Y-27632 (Tocris Bioscience, Minneapolis, Minnesota, USA) into Aggrewell 800 microwell plates (Stemcell Technologies) at a density of 3×10^6 cells/well. On day -1, cells were given a half volume media change of mTeSR Plus. On day 0, embryoid bodies were transferred to ultra-low attachment 10 cm dishes (Corning, Corning, New York, USA) on an orbital shaker, and the media was changed to neural induction media (NIM) comprised of DMEM/F-12, 20% knock-out serum replacement, 1X GlutaMAX, 1X MEM non-essential amino acids, 100 U/ml penicillin, 100 μg/ml streptomycin, 100 μM 2-mercaptoethanol (Thermo Fisher Scientific) with 5 μM Dorsomorphin (Tocris), and 10 μM SB-431542 (Cayman Chemical, Ann Arbor, Michigan, USA) (NIM+SB/DM media). NIM+SB/DM media was changed daily from days 1 to 3. On day 4 organoids were split into two batches with half undergoing a dorsalization protocol and half undergoing a ventralization protocol. For dorsalization, organoids were given NIM+SB/DM media daily until day 6 when they were given NM+EGF/FGF media comprised of neural media (NM, Neurobasal-A, 1X B27 supplement without vitamin A, 1X GlutaMAX, 100 U/ml penicillin, 100 μg/ml streptomycin) with 20 ng/ml EGF (R&D Systems) and 20 ng/ml FGF2 (Thermo Fisher/Peprotech). NM+EGF/FGF media was changed daily from days 7 to 14, then changed every other day from days 15 to 23. Starting on day 25 organoids were switched to NM with 20 ng/ml BDNF (Thermo Fisher/Peprotech) and 20 ng/ml NT3 (Thermo Fisher/Peprotech) and media changed every third day. After day 46 organoids were switched to NM media, changed every third day.

For ventralization, organoids were given NIM+SB/DM/IWP media, comprised of neural induction media with 5 μM Dorsomorphin (Tocris), 10 μM SB-431542, and 5 uM IWP-2 (Selleckchem, Houston, Texas, USA) on days 4 and 5. On day 6 they were switched to NM+EGF/FGF/IWP, comprised of neural media with 20 ng/ml EGF, 20 ng/ml FGF2, and 5 uM IWP-2. NM+EGF/FGF/IWP media was changed daily from days 7 to 11 and NM+EGF/FGF/IWP/SAG (Selleckchem, 100 nM)/RA (Sigma Aldrich, St. Louis, Missouri, USA, 100 nM) media was given daily on days 12 to 14. NM+EGF/FGF/IWP/SAG/AlloP (Cayman, 100 nM) media was then given every other day from days 15 to 23. Starting on day 25 organoids were switched to NM with 20 ng/ml BDNF and 20 ng/ml NT3, changed every third day. After day 46 organoids were switched to NM media, changed every third day. To keep track of the gross morphology and size of organoids, 5x bright-field images of organoids at day 5 of differentiation were captured using an Olympus Infinity microscope every other day until day 21, then weekly until day 35.

#### Western blot

Organoids were collected into 1.5 ml Eppendorf tubes and media removed. No extra washes were done to avoid activating mTOR signaling. NP-40 lysis buffer (40 mM HEPES, pH 7.5, 120 mM NaCl, 1 mM EDTA, 1% Triton X-100, 1% NP-40) with freshly added protease inhibitor cocktail (Roche, Basel, Switzerland, #14132300) and phosphatase inhibitor cocktail (Thermo Fisher #A32957) was added to the tubes and samples sonicated until no pellet remained. Protein lysate samples were centrifuged at 14,000 rpm for 30 min at 4°C and the supernatant transferred to new tubes. Protein quantitation was carried out using a Qubit fluorometer and the Qubit Protein Assay Kit (Thermo Fisher).

Protein samples were prepared by boiling at 95 °C for 5 min in SDS sample buffer and subsequently separated on 4-15% Tris-Glycine precast gels (Bio-Rad, CA, USA). Following electrophoresis, proteins were transferred to nitrocellulose membranes using a semi-dry transfer system (Bio-Rad). Membranes were blocked at room temperature for 2 hours in Intercept Tris-buffered saline (TBS) blocking buffer (Li-COR, NE, USA), then incubated overnight at 4 °C with primary antibodies diluted in TBS blocking buffer with 0.1% Tween-20 (TBS-T). After three washes with TBS-T, membranes were incubated with IRDye-conjugated secondary antibodies (Li-COR) for 2 hours at room temperature in the dark. Immunoreactive bands were visualized using the ChemiDoc MP imaging system (Bio-Rad).

Band intensities were quantified using Image Lab software (Bio-Rad). STRADA expression was normalized to the housekeeping protein GAPDH to control for loading variability. For phospho-protein analysis, the signal intensity of phosphorylated proteins (pS6 or p4EBP1) was normalized to the total corresponding protein (S6 or 4EBP1) within each sample. Normalized values were used for statistical comparisons between groups.

#### RNA isolation and RT-qPCR

Total RNA was isolated with Quick-RNA Miniprep Kit (Zymo, Irvine, California, USA) according to the manufacturer’s instructions. cDNA was synthesized with the SuperScript™ III First-Strand Synthesis SuperMix kit (Thermo Fisher). RT-qPCR was carried out with Power SYBR Green PCR Master Mix (Thermo Fisher, 4367659). Fold changes were calculated using the 2^-ΔΔCt method: ΔΔCt = [(Ct gene of interest - Ct internal control) KO - (Ct gene of interest - Ct internal control) WT], with GAPDH being an internal control. Primer sets to check STRADA and GAPDH mRNA levels are in the key resources table.

#### EdU-labeling/quantification/analysis

For cell proliferation index and cell cycle exit ratio analysis, organoids were treated at day 14 with a final concentration of 10 µM EdU for 2 h or 24 h, respectively, before harvesting for fixation. Following fixation, cryosectioning, and immunostaining (co-stained with KI-67), EdU incorporation was detected using the Click-iT EdU Cell Proliferation Kit (Thermo Fisher Scientific) following the manufacturer’s protocol. Proliferation index was calculated as EdU+/all cells. Cell cycle exit ratio was calculated as EdU+ Ki67-/all EdU+ cells.^30^

For cell fate analysis, organoids were treated with 10 µM EdU at day 32 and harvested at day 35. EdU-labeled cells were visualized using the same protocol as above and co-stained with lineage-specific markers, including PAX6 (progenitors)/ EOMES (intermediate progenitors)/ BCL11B (neurons) for dorsal and SOX2 (neural progenitors)/NeuN (neurons) for ventral. Quantification was performed by calculating the percentage of EdU+ cells co-expressing each marker.

#### Rapamycin treatment

For rapamycin treatment, a portion of STRADA KO dorsal and ventral organoids were separated out and treated with rapamycin at a final concentration of 20 nM starting at day 12 of differentiation and maintained in the media till collection. The remaining portion of STRADA KO, together with the control organoids, were treated with an equal volume of DMSO added to the media starting from day 12.

#### Immunohistochemistry

Cortical organoids were fixed in 4% paraformaldehyde for 20–30 min at 4°C, cryoprotected overnight with 30% sucrose in phosphate-buffered saline, and then embedded in Tissue-Tek OCT embedding medium (Sakura Finetek, Torrance, California, USA) and stored at −80°C. A Leica CM1850 cryostat was used to generate 20 μm thick sections that were collected on Superfrost Plus glass slides (Thermo Fisher) and stored at −20°C. For immunostaining, the sections were outlined with a lipid pen and then washed 3 times with phosphate-buffered saline (PBS) for 5 min each. The sections were permeabilized with PBS with 0.2% Triton-X100 for 20 min and then incubated with blocking buffer (PBS with 0.05% Tween-20, 5% normal goat serum, and 1% bovine serum albumin) for 1 hr at room temperature (RT). The slides were incubated with primary antibodies diluted in blocking buffer overnight at 4°C in a humidified chamber. The slides were washed three times with PBS with 0.05% Tween-20 (PBS-T) for 10 min at RT, and incubated with AlexaFluor-conjugated secondary antibodies (Invitrogen, Carlsbad, California, USA) for 1–2 hr at RT. The slides were then washed once with PBS-T for 10 min at RT and then incubated in 2 μg/ml bis-benzimide diluted in PBS for 5 min to label nuclei. The slides were washed three times with PBS-T for 10 min at RT, and mounted with Glycergel mounting medium (Agilent Dako, Santa Clara, California, USA). 20x images were obtained on an Andor BC43 spinning-disc confocal microscope using the Fusion BC43 acquisition software (Oxford Instruments, England).

#### Image analysis

Images were analyzed using Fiji (NIH, Bethesda, MD, USA).^71^ All image quantifications were performed automatically using custom-written Fiji macros. For organoid size measurement, auto-thresholding was applied to contrast-enhanced bright-field images to distinguish organoid structures from the background. The images were then converted to binary masks, followed by particle analysis to measure area and perimeter.

For mean fluorescence intensity (MFI) quantitation, Z-Stacks were initially converted to single plane images by selecting the z position with the highest mean fluorescence. Using a merged single-plane image, a mask was created representing the total area of the organoid, excluding any cracks or holes in the section. The mean fluorescence intensity was then measured separately for each fluorophore within the mask. Custom thresholds were established for each channel based on signal intensity, and identical thresholding settings were consistently applied to each corresponding channel across all samples within the same immunostaining condition. Each marker was evaluated across at least three independent biological replicates. A minimum of four organoids per cell line were analyzed per replicate to ensure reproducibility.

To perform cell counting for EdU-labeled lineage analysis and interneuron specification, thresholding was applied to convert the images to binary masks for each channel. Particles were counted using the “Analyze Particles” function, in which the parameters were set to exclude small particles (<10 pixels) and noise. To identify double positive cells, an image calculator was used to generate a binary mask representing the co-localization of two markers, followed by particle analysis. The numbers of each single channel and double-positive cells were recorded for each trial. Quantification was performed across at least three independent biological replicates, with analysis of at least four organoids per cell line per replicate.

#### Single Cell RNA Sequencing

##### Sample Preparation

Day 35 or day 77 organoids were dissociated using the Worthington papain dissociation kit. dhCOs and vhCOs were collected in parallel from the same differentiation for each timepoint. Two mL of papain/DNase solution was used for between 5 to 20 organoids, placed in a well of a low-adherence 24-well plate, depending on size of organoids for each respective condition. At least 5 organoids were pooled for each dissociation. Organoids were incubated at 37 ºC for 45 minutes with gentle orbital shaking followed by 10 hard passes through a P1000 micropipette. Organoids were incubated again for 15 minutes followed by 5 additional passes through a P1000 tip before the cell solution was filtered through a 40 µm cell strainer. Cells were collected by centrifugation and cell count and viability assessed before proceeding with sample-specific hashtag oligonucleotide-conjugated antibody labeling (TotalSeq-B, Biolegend) following the manufacturer’s protocol. This allowed pooled analyses of multiple (up to 6) samples to reduce cost and batch effects, with subsequent de-multiplexing of scRNA-seq data based on the hashtags.

For all samples from differentiations treated with rapamycin, labeled cells from all dorsal samples and all ventral samples were then separately mixed in equal quantities to generate one combined dorsal sample and one combined ventral sample from each timepoint before submitting to the University of Michigan Advanced Genomics Core. A separate day 35 sample of dhCOs and vhCOs generated from only lines C1 and M1 without rapamycin treatment was also prepared for scRNA-Seq where equal cell numbers from each of the four samples were combined before submission for sequencing. Cell viability was found to be greater than 72% in all samples by fluorescent viability assay.

##### Library Preparation and Sequencing

Single-cell suspensions were processed using the 10X Genomics Chromium platform, and scRNA-seq libraries were sequenced according to the manufacturer’s protocol (Illumina NovaSeqXPlus). Each paired-end read contains a cell barcode and a UMI in Read 1, and the transcript-specific read in Read 2 with a read length of 151 nt. Separately, each cell generates a hashtag oligo count vector for the hashtags used, allowing the cells to be assigned to the organoid samples they come from.

##### Data processing and filtering

BCL Convert Software v4.0 (Illumina) was used to generate de-multiplexed Fastq files from BCL files together with barcode information. Transcript reads were analyzed using 10x Genomics Cell Ranger (v7.2.0) with GRCh38-2020-A as the reference genome, to obtain single-cell gene expression count matrices. For quality control, we excluded cells predicted to be multiplets by the Cell Ranger pipeline, as well as cells that had <1000 detected genes and >10% mitochondrial transcripts.

##### Clustering, visualization, and cluster annotation

We performed cell clustering separately on the dorsal and the ventral samples, with all time points combined. Raw transcript reads were normalized following the SCTransform normalization workflow from Seurat package (v5.0.1, https://satijalab.org/seurat/).^72–76^ Dimensionality reduction and unsupervised cell clustering were also done using Seurat package (v5.0.1), to generate UMAPs. Initially automated cell type annotation of the clusters was performed using SingleR (v1.16.0) (https://bioconductor.org/books/release/SingleRBook/)^77^ with a reference data set,^78^ and then refinement of the assigned clusters was done manually using known markers for cell types from the brain.^79^ Similar clusters were combined, and clusters containing sick/dying cells, marked by high expression of mitochondrially-encoded genes, and mesenchymal cells were removed.

##### Differentially expressed genes and pathways

Differentially expressed genes were determined by using FindAllMarkers function from Seurat package, with a minimum log2 fold change of 0.25 and an adjusted *p*-value of less than 0.05. Reactome pathway enrichment analysis for each group of pairwise comparison was performed by using DAVID Bioinformatic platform.^36,37^ Enrichment score was calculated as -log_10_ of the geometric mean of the p-values for the pathways.

Clusters of pathways were determined for the following dhCO cell type clusters: Cortical radial glia, dividing; cortical radial glia, non-dividing; cortical radial glial and outer radial glia, non-dividing; cortical intermediate progenitors; and cortical excitatory neurons. For the vhCOs, pathway cluster analysis was performed for the following cell type clusters: Ganglionic eminence radial glia, dividing; ganglionic eminence radial glia, non-dividing; ventral glial progenitors and outer radial glia; ganglionic eminence intermediate progenitors; medial ganglionic eminence pallial neurons; medial ganglionic eminence pallial neurons, metabolically active; and subpallial medial ganglionic eminence interneurons.

### QUANTIFICATION AND STATISTICAL ANALYSIS

Data were organized and analyzed in EXCEL (Microsoft, Seattle, WA, USA). Graphing and statistical analyses were performed using Prism 10 (Graphpad, La Jolla, CA, USA). Size measurement, cell counts, qPCR results, and MFI quantitation were tested for statistically significant differences. Comparisons between two groups were performed using the non-parametric Mann-Whitney test, while comparisons involving more than two conditions were conducted using the one-way ANOVA followed by Dunn’s multiple comparison test or non-parametric Kruskal Wallis test with multiple comparisons. Results are presented as mean ± SEM.

For organoid size analysis in Fig. 1B, we used SAS (SAS Institute, Inc., Cary, NC) and modeled the data using mixed models of individual growth, where an “individual” was considered a region (dorsal or ventral) for a given cell line (e.g. C1, C2, M1, M2), with or without rapamycin, in a given differentiation. These individuals were considered nested within experiments and not assumed to be independent from other individuals within that same experiment for the purpose of statistical modeling. We used separate models for the dorsal and ventral region data. Initial graphic displays of the data suggested that the pattern of growth was non-linear, therefore, we tested a quadratic term for time in the mixed model. This quadratic term was not significant in the dorsal model but was significant in the ventral model; therefore, the quadratic term was retained in the ventral model but not the dorsal model. Mixed models were run using SAS Proc Mixed with random effects for intercepts and time for measurement unit nested within experiment (variance components covariance structure), using maximum likelihood estimation and Kenward-Roger degree of freedom adjustments.

