## Supplemental Figures and Table for "Delayed forebrain excitatory and inhibitory neurogenesis in STRADA-related megalencephaly via mTOR hyperactivity"

### SUPPLEMENTARY INFORMATION

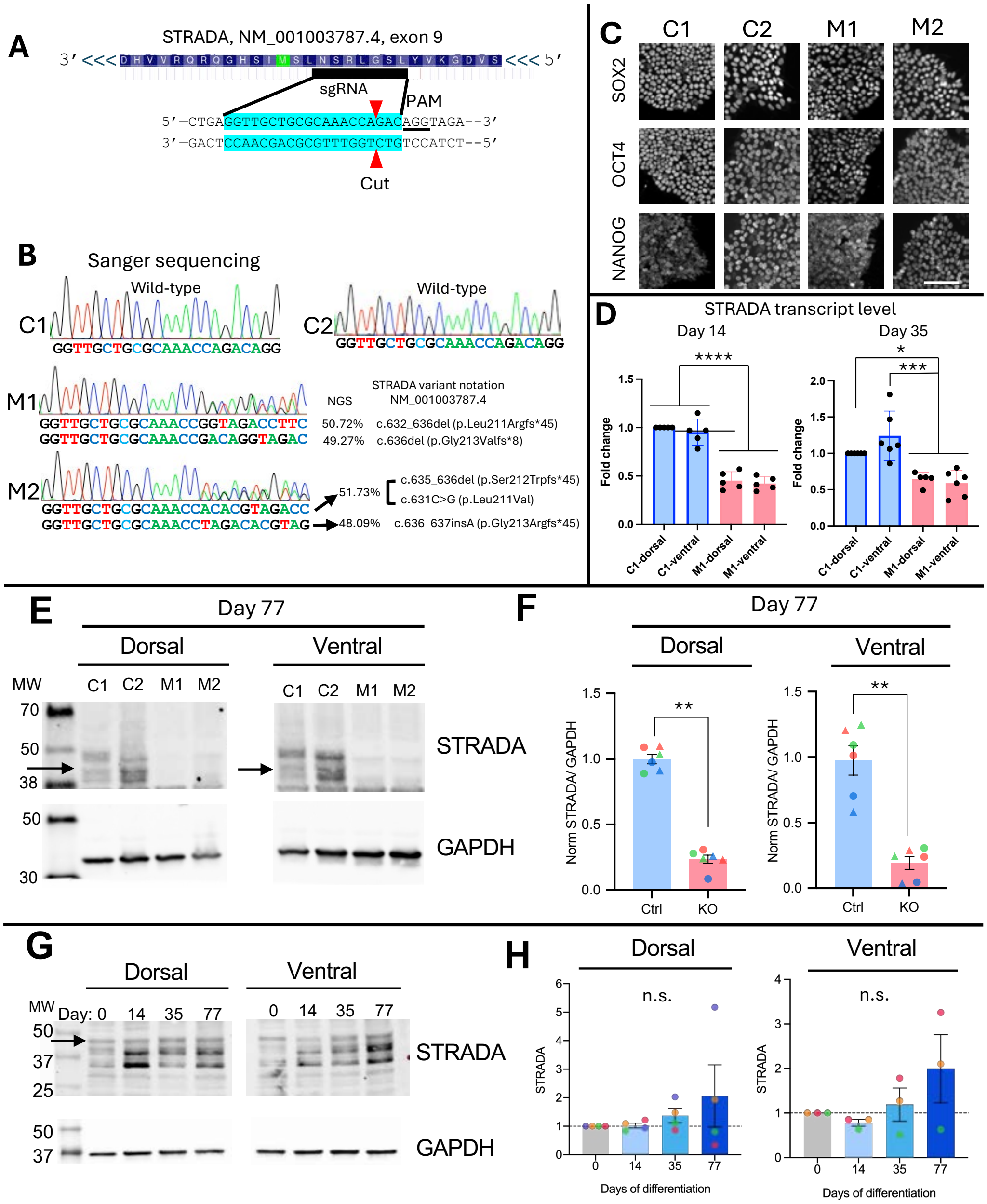


**Figure S1: Generation and characterization of STRADA KO iPSC lines and organoids. (A) CRISPR-editing strategy for generating STRADA KO lines.** A guide RNA (sgRNA) targeting exon 9 of the human *STRADA* gene introduced loss of function mutation. **(B)** Sanger and next-generation sequencing analysis of C1, C2, M1, and M2 iPSC lines. **(C)** iPSC pluripotency verification. Immunostaining confirmed expression of SOX2, OCT4, and NANOG across all lines. Scale bars, 100 µm. **(D)** qRT-PCR results for *STRADA* mRNA expression in day 14 and 35 C1 and M1 dhCOs and vhCOs. **(E)** Western blotting results for STRADA protein expression in day 77 control and STRADA KO dhCOs and vhCOs. The black arrow indicates the target band location of STRADA. **(F)** Western blot quantification from E. **(G)** Western blot for STRADA protein expression in a control line (C1) over the course of dhCO and vhCO differentiation. **(H)** Quantification of STRADA Western blotting results from G. Quantification dots represent biological replicates from 3-6 batches of differentiations per condition, with colors indicating differentiations and shapes indicating genotypes (circles for C1/M1, triangles for C2/M2). Mean ± SEM is displayed, with statistical significance determined by Mann-Whitney test, $^{**}p<0.01$, $^{***}p<0.001$, $^{****}p<0.0001$.

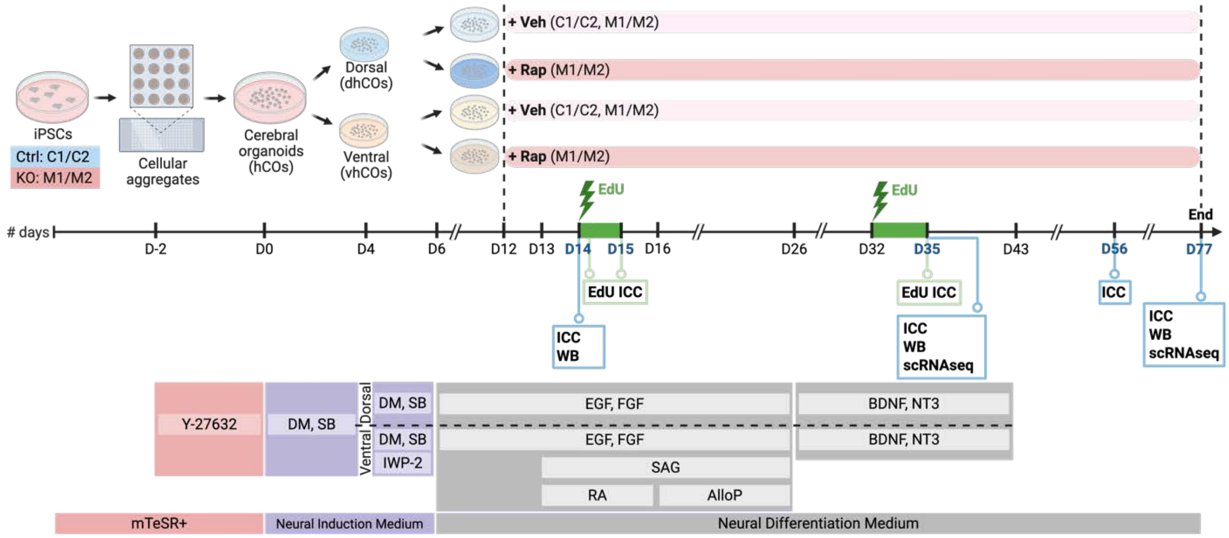


**Fig. S2**: **Study design and schematic of the workflow.** ICC, immunohistochemistry. WB, Western blotting. scRNAseq, single cell RNA-sequencing. Created with *Biorender.com.* Rap: rapamycin, Veh: vehicle.

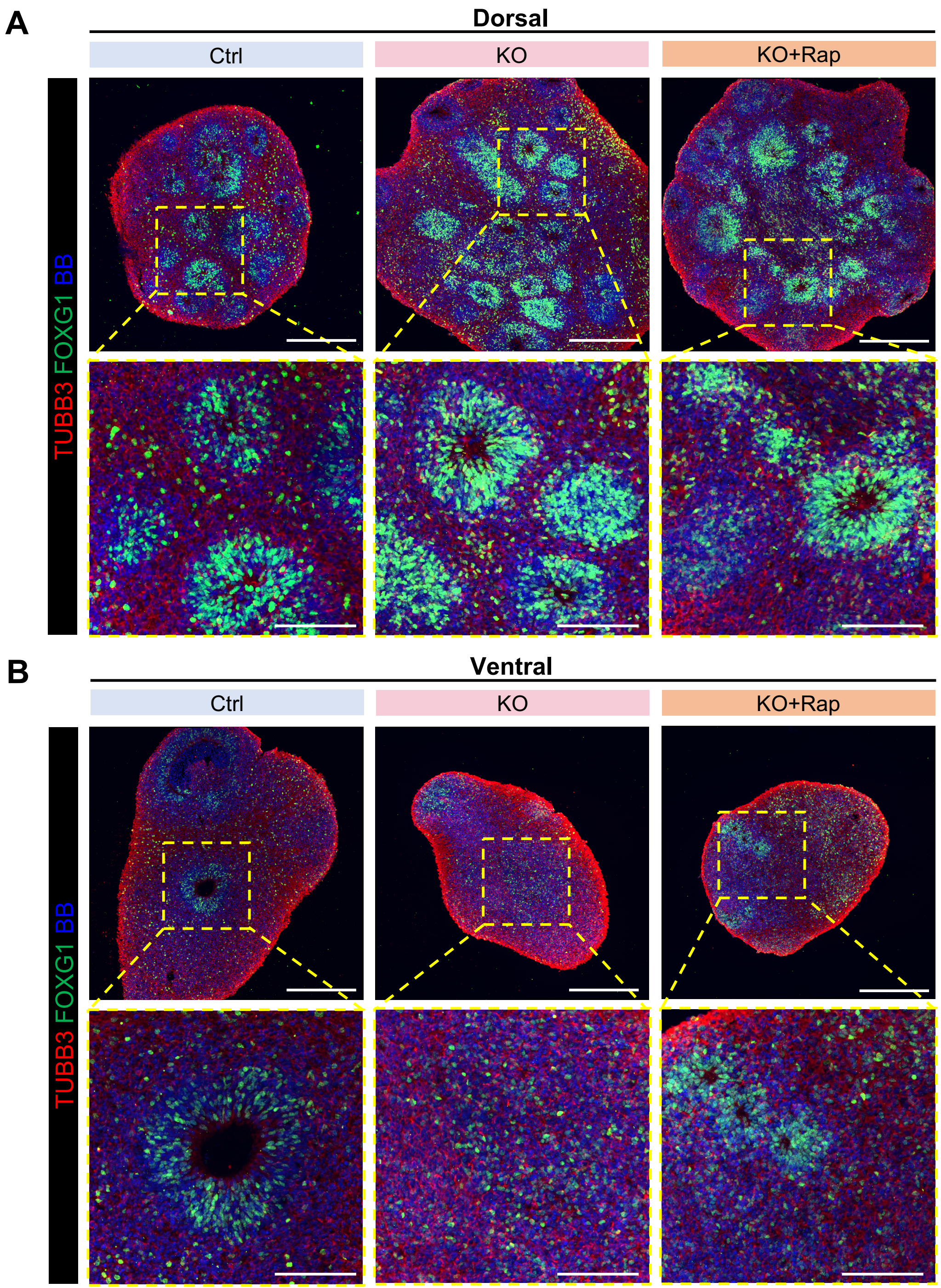


**Fig. S3: dhCO and vhCO differentiation protocols generate cortical organoids with forebrain identity.** Representative images for immunostaining for TUBB3 and FOXG1 in day 56 control and STRADA KO dhCOs (**A**) and vhCOs (**B**) treated with vehicle or rapamycin. Scale bars, 300 μm (100 μm for zoomed-in image).

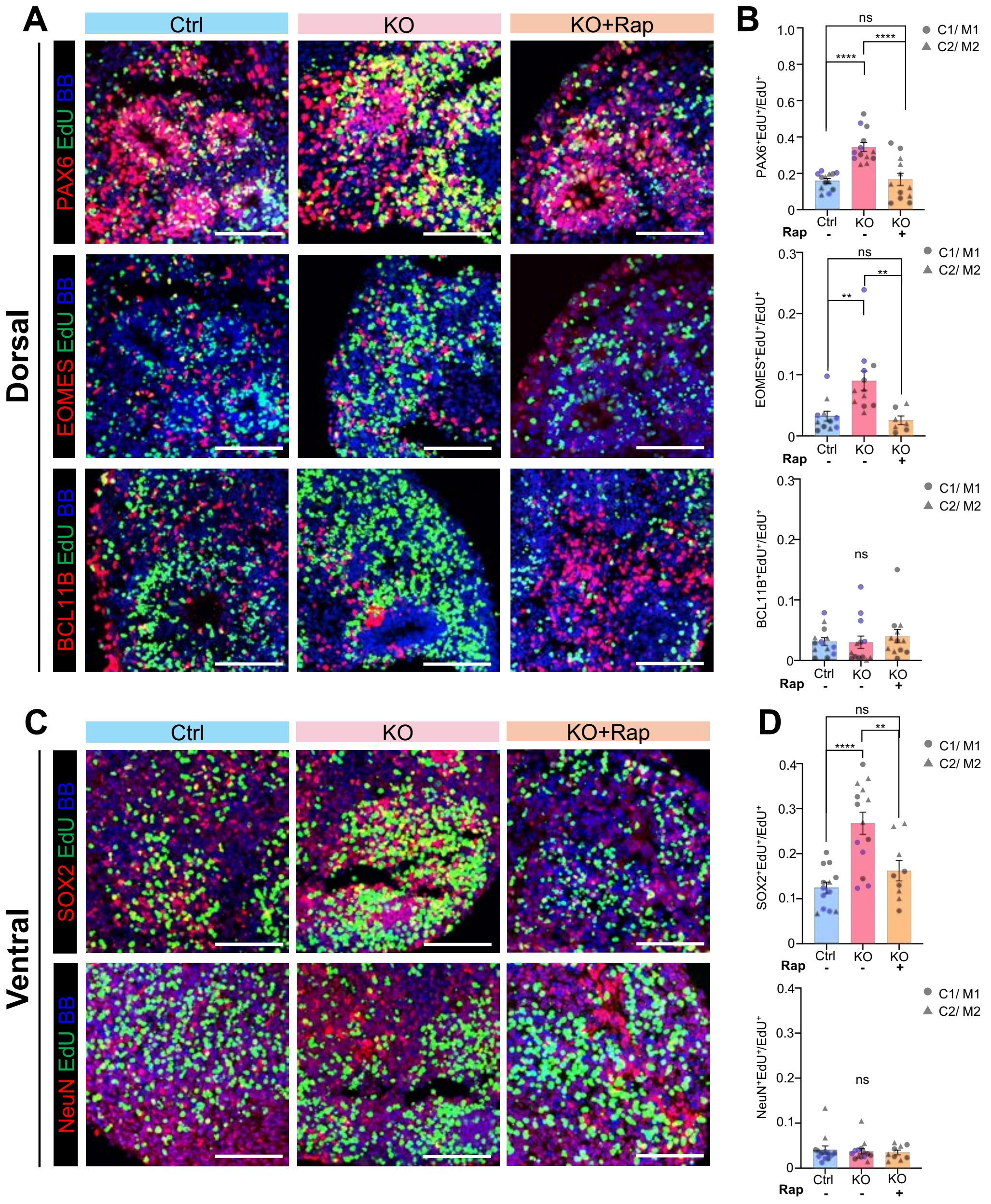


**Fig. S4: STRADA KO dhCOs and vhCOs show increased ratios of cycling neural progenitors. (A)** Immunostaining for Pax6, EOMES, and BCL11B in day 35 control and KO dhCOs, treated with either vehicle or rapamycin. Organoids were pulse labeled with EdU from day 32. **(B)** Quantitation of progenitors ($PAX6^{+}EdU^{+}/EdU^{+}$), intermediate progenitors ($EOMES^{+}EdU^{+}/EdU^{+}$), and neurons ($BCL11B^{+}EdU^{+}/EdU^{+}$) in dorsal organoids from panel A. **(C)** Immunostaining for SOX2 and NeuN in day 35 control and KO vhCOs, treated with vehicle or rapamycin from day 12. EdU was administered from day 32. **(D)** Quantification of neural progenitors ($SOX2^{+}EdU^{+}/EdU^{+}$) and neurons ($NeuN^{+}EdU^{+}/EdU^{+}$) in vhCOs from panel C. For all quantifications, lineage ratios were calculated as the percentage of EdU+ cells. Each symbol represents an individual organoid, compiled from various batches of differentiations (color-coded). Data are presented as mean ± SEM. Statistical significance was determined by one-way ANOVA. *$p<0.05$, **$p<0.01$, ***$p<0.001$, ****$p<0.0001$. Scale bars, 100 μm.

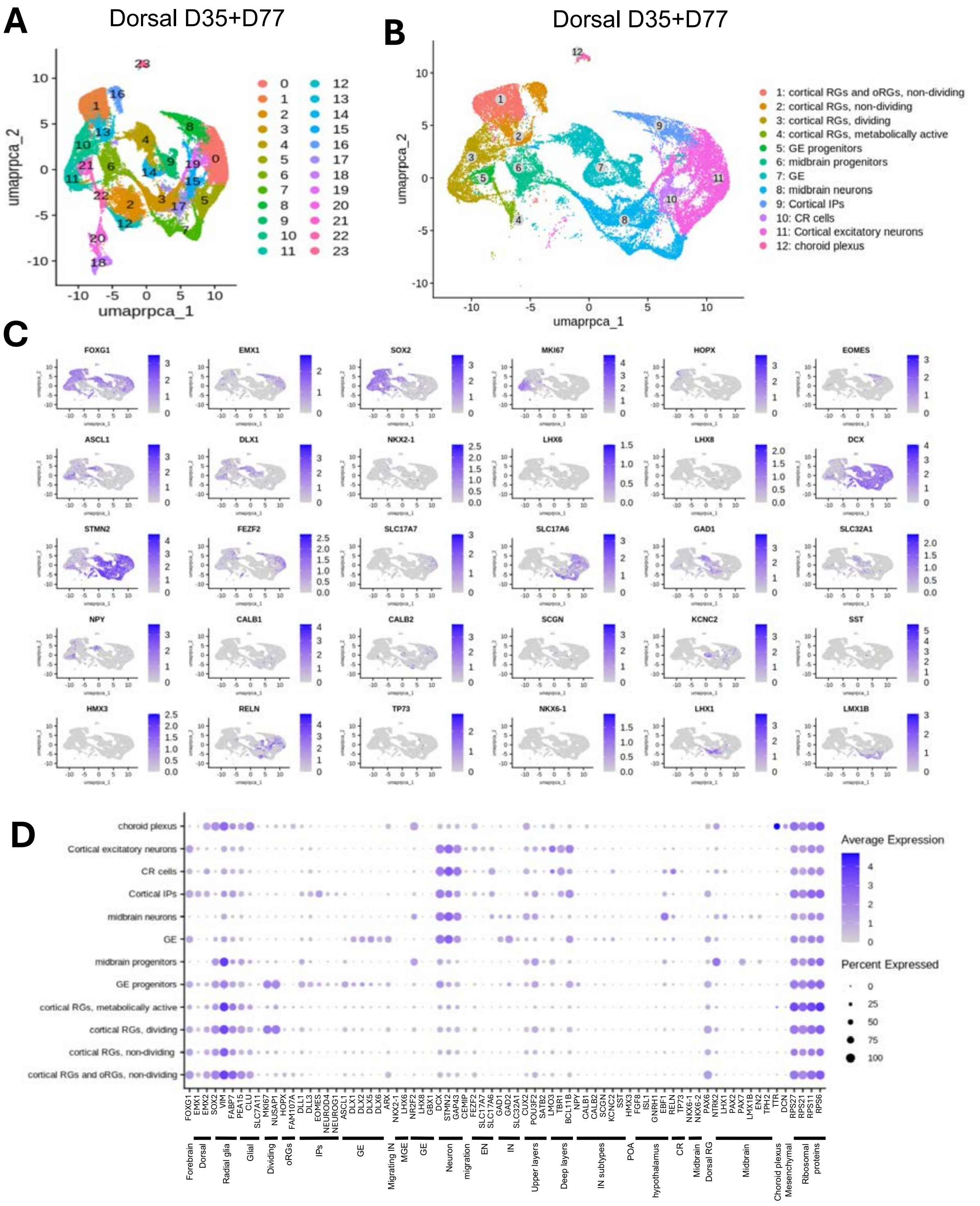


**Fig S5: Cell type clustering analysis of scRNA-Seq results from control and STRADA KO day 35 and day 77 dhCOs treated with vehicle or chronic rapamycin**. **(A)** Combined UMAP plot of all dhCO samples with cluster identification. Legend at right displays the cell type identity for each cluster. **(B)** Feature plots of cell type specific markers and selected genes in control and STRADA KO dhCOs. **(C)** Dot matrices of marker gene expression levels in each cluster. Scale bars for relative expression level and percentage of cells within the cluster expressing the marker are included at the right of each panel.

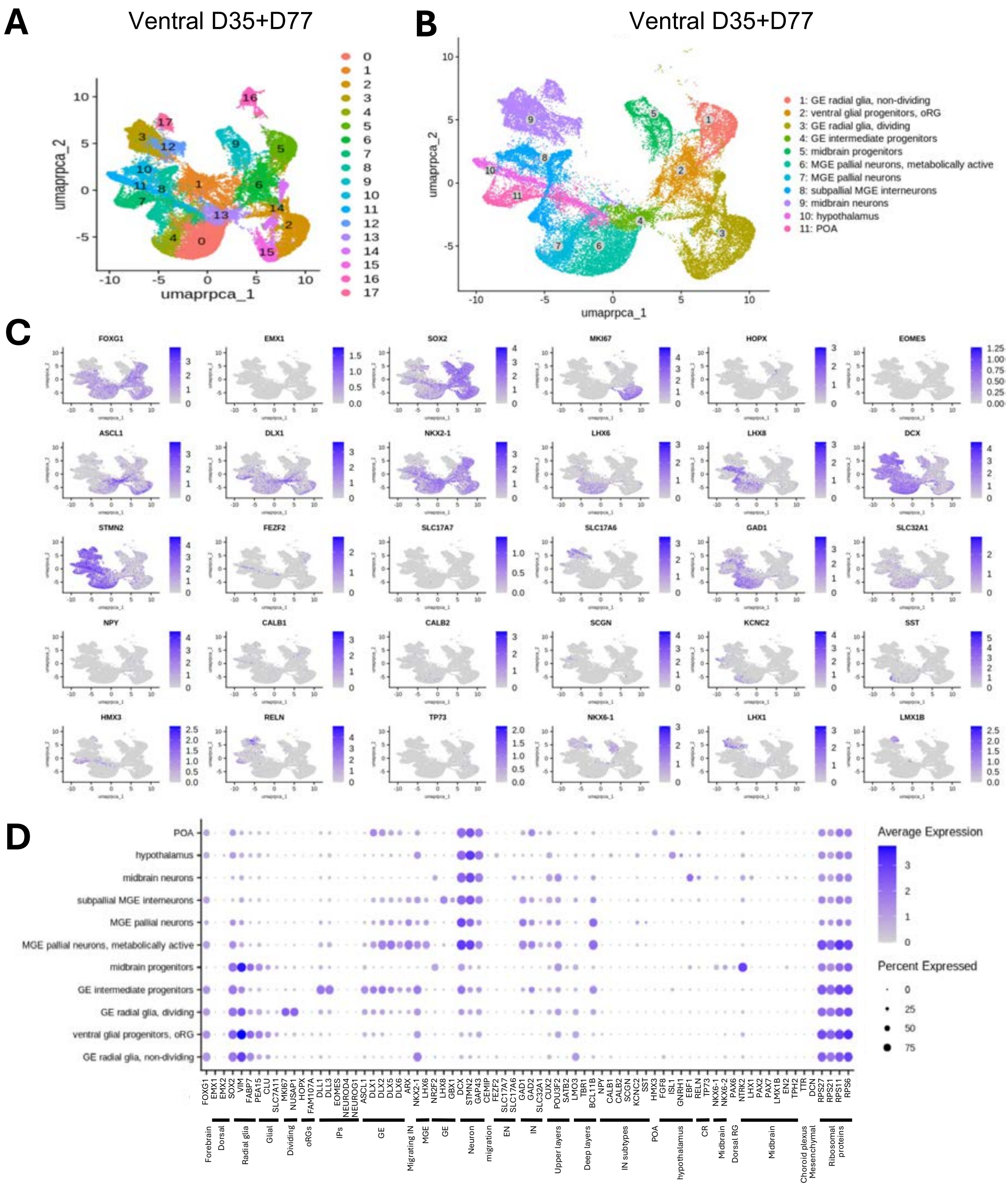


**Fig. S6: Cell type clustering analysis of scRNA-Seq results from control and STRADA KO day 35 and day 77 vhCOs treated with vehicle or chronic rapamycin. (A)** Combined UMAP plot of all vhCO samples with cluster identification. Legend at right displays the cell type identity for each cluster. **(B)** Feature plots of cell type specific markers and selected genes in control and STRADA KO vhCOs. **(C)** Dot matrices of marker gene expression levels in each cluster. Scale bars for relative expression level and percentage of cells within the cluster expressing the marker are included at the right of each panel.

**Fig. S7**

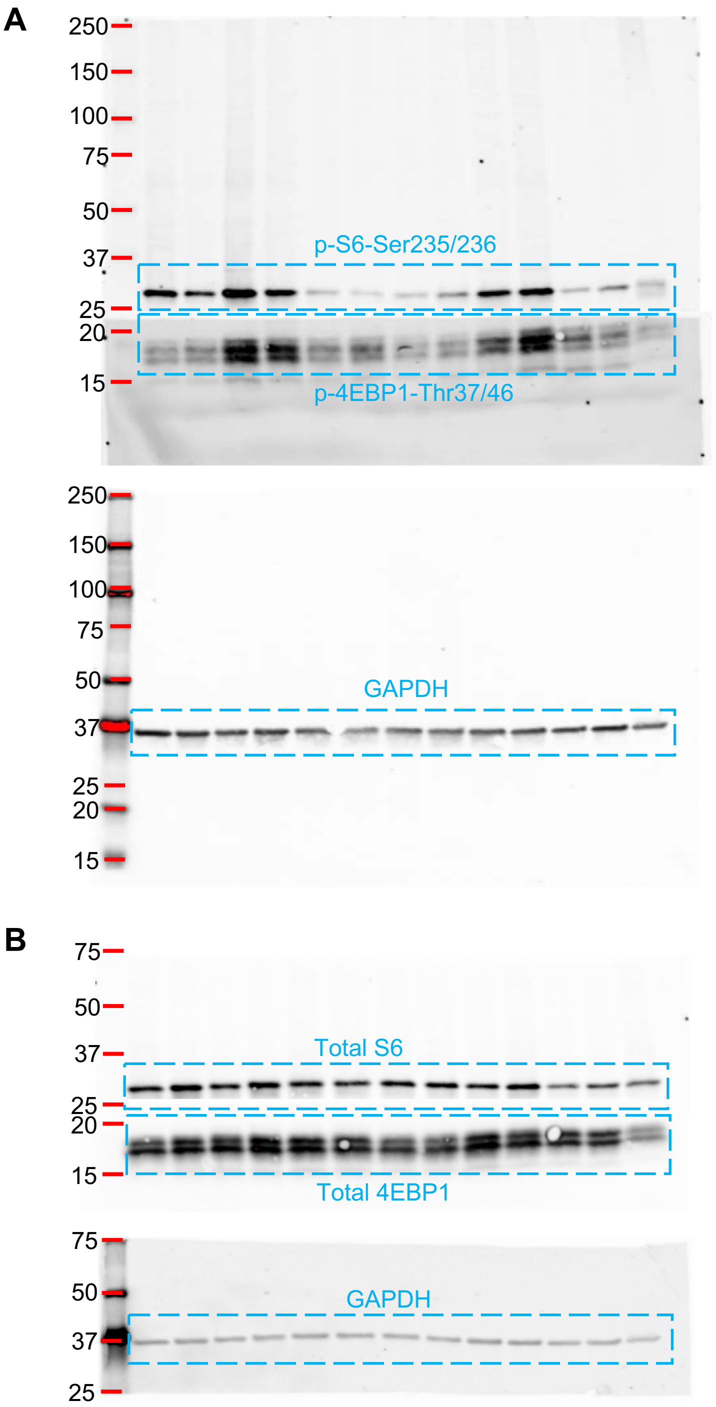
A
250
150
100
75
50
37
25
20
15
p-S6-Ser235/236
p-4EBP1-Thr37/46
GAPDH
B
Total S6
Total 4EBP1
GAPDH

**Fig. S7, continued**

C

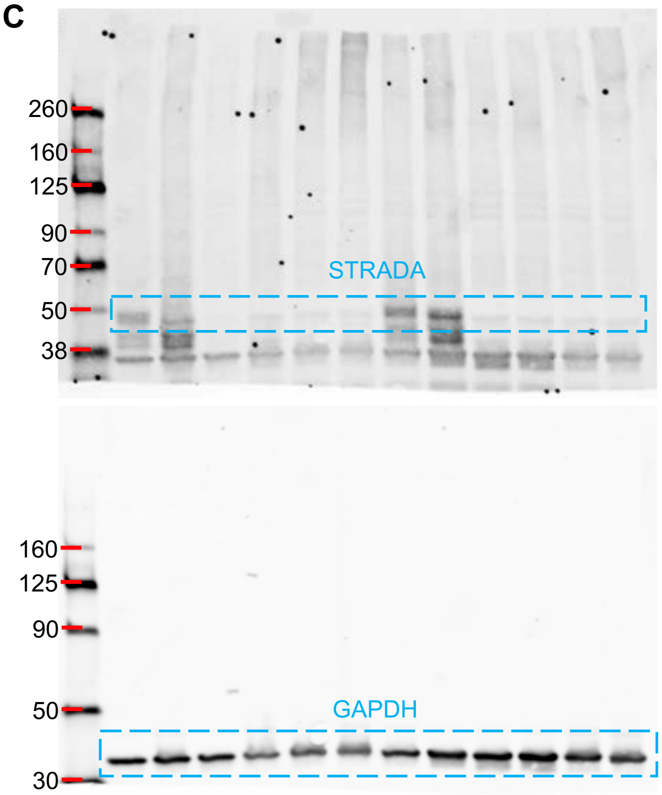


D

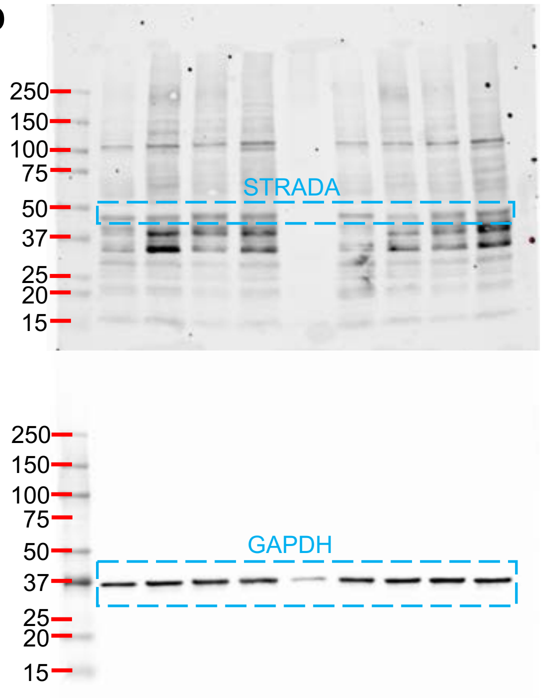


**Fig. S7: Uncropped western blot images, referring to Fig. 2A and 2C and S2 (A**) For phosphorylated S6, phosphorylated 4EBP1, and **(B)** for total S6 and total 4EBP1, each with GAPDH as loading control. **(C)** STRADA expression, referring to Fig S2E. **(D)** STRADA expression, referring to S2G.

**Table S1. Single cell RNA-sequencing samples**

| Multiplexed samples | Days in vitro | Regionalization | |
|---|---|---|---|
| | | Dorsal | Ventral |
| Sample 1 | 35 | C1, M1 | C1, M1 |
| Sample 2 (dorsal) + Sample 3 (ventral) | 35 | C1, C2, M1, M2, M1 with rapa, M2 with rapa | C1, C2, M1, M2, M1 with rapa, M2 with rapa |
| Sample 4 (dorsal) + Sample 5 (ventral) | 77 | C1, C2, M1, M2, M1 with rapa, M2 with rapa | C1, C2, M1, M2, M1 with rapa, M2 with rapa |

Five samples were submitted for single cell RNA-sequencing, with sample 1 containing 4 multiplexed samples, and samples 2-5 each containing 6 multiplexed samples for a total of 28 samples. C1 and C2 refer to two control cell lines, and M1 and M2 refer to two STRADA knockout cell lines.
